# Cryo-EM structures of ρ1 GABA_A_ receptors with antagonist and agonist drugs

**DOI:** 10.1101/2024.07.30.605774

**Authors:** Chen Fan, John Cowgill, Rebecca J. Howard, Erik Lindahl

## Abstract

The family of ρ-type GABA_A_ receptors includes potential therapeutic targets in several neurological conditions, and features distinctive pharmacology compared to other subtypes. Here we combine structures, recordings and simulations to characterize the binding and conformational impact of the drugs THIP (a non-opioid analgesic), CGP36742 (phosphinic acid inhibitor) and GABOB (an anticonvulsant) on a human ρ1 GABA_A_ receptor. We identify a distinctive binding pose of THIP in ρ1 versus neuronal α4β3δ GABA_A_ receptors, offering a rationale for its inverse effects on these subtypes. CGP36742 binding is similar to the canonical ρ-type inhibitor TPMPA, supporting a shared mechanism of action among phosphinic acid inhibitors. Binding of GABOB is similar to that of GABA, but produces a mixture of primed and desensitized states, likely underlying its weaker agonist activity. Together, these results elucidate interactions of a ρ-type GABA_A_ receptor with therapeutic drugs, offering mechanistic insights and a prospective basis for further pharmaceutical development.

## Introduction

γ-Aminobutyric acid (GABA) is the major inhibitory neurotransmitter in the human central nervous system and it is crucial for the balance of neuronal excitation and inhibition^1^. Two types of GABA receptors have been identified based on their mechanism of action: GABA_A_ receptors are GABA-gated chloride ion channels, while GABA_B_ receptors are G protein-coupled receptors^1^. GABA_A_ receptors belong to the pentameric ligand-gated ion channel (pLGIC) superfamily, which also contains ionotropic receptors for acetylcholine, glycine, and serotonin^2^. Channels in this family share a common architecture, with each of the five subunits contributing to an extracellular domain (ECD) with 10 strands (β1–β10) and a transmembrane domain (TMD) with 4 helices (M1–M4). The orthosteric ligand binding site is located at the subunit interface in the ECD, composed of loops A, B and C from the principal subunit and D, E and F from the complementary subunit. Ligand binding is thought to induce an anticlockwise rotation of the ECD relative to the TMD when viewed from the extracellular side. These conformational changes propagate to the TMD, causing the opening of the pore formed by the M2 helices along the central axis, to allow ions to transit the lipid bilayer^3^. In contrast, GABA_B_ receptors mediate relatively slow and prolonged synaptic inhibition, with presynaptic GABA_B_ receptors suppressing neurotransmitter release, and postsynaptic GABA_B_ receptors causing hyperpolarization of neurons^4^.

In humans, GABA_A_ receptors are homo- or hetero-pentamers formed from a selection of 19 different subunits (α1-6, β1-3, γ1-3, ρ1-3, δ, ε, π and θ). Channels formed from ρ subunits were previously named GABA_C_ receptors due to their distinctive physiological and pharmacological properties, including higher sensitivity to but slower activation by GABA relative to classical synaptic and extrasynaptic subtypes; this subtype is also relatively insensitive to bicuculline, barbiturates and benzodiazepines^5^, but sensitive to phosphinic acid compounds including (1,2,5,6-tetrahydropyridin-4-yl)methylphosphinic acid (TPMPA)^6^, which provides possibilities to selectively modulate particular subforms of human GABA_A_ receptors. Of the 3 types of ρ subunits found in mammals, ρ1 is expressed particularly in the retina, while ρ2 and ρ3 are widely distributed in brain^7, 8^. ρ-type receptors play important roles in physiological processes including visual transduction^9^, postnatal neurodevelopment^10^, pain sensation^11^, and sleep-wake cycles^12^, and are potential therapeutic targets in myopia, sleep disorders, learning and memory disruption, peripheral nociception and anxiety^13, 14^.

Several drugs act on ρ-type GABA_A_ receptors, including the synthetic agents 4,5,6,7-tetrahydroisoxazolo[5,4-c]pyridin-3-ol (THIP, also known as gaboxadol) and 3-aminopropyl-*n*-butylphosphinic acid (CGP36742, also known as SGS742) and the natural product γ-amino-β-hydroxybutyric acid (GABOB, also known as buxamine) (Supplementary Fig.1). THIP, a conformationally constrained derivative of muscimol, was developed as a non-opioid analgesic and antinociceptive agent^15, 16^, and was a candidate in clinical trials for the treatment of insomnia^17^, Fragile X syndrome^18^ and Angelman syndrome^19^. CGP36742 was the first GABA_B_ receptor antagonist in clinical trials^20^, but is also an orally active antagonist of ρ-type GABA_A_ receptors^21, 22^, and has shown therapeutic potential for the treatment of cognitive deficits^23, 24^. As a metabolite of GABA, GABOB is found endogenously in the mammalian central nervous system, but it is also an anticonvulsant used in the treatment of epilepsy^25^. The hydroxyl group at the C3 position of GABOB generates a stereogenic center, resulting in R and S enantiomeric forms; although (R)-GABOB is a modestly more potent anticonvulsant, the compound is applied clinically as a racemic mixture^26^. Despite their therapeutic relevance, the structural foundations of these drugs’ selectivity and other functional properties at ρ-type GABA_A_ receptors remain unclear.

Here, combining structures, electrophysiology and molecular dynamics (MD) simulations, we characterize the binding and structural impact of THIP, CGP36742 and GABOB on human ρ1 GABA_A_ receptors. We identify a distinctive binding pose of THIP in ρ1 versus the extrasynaptic neuronal α4β3δ GABA_A_ receptors, offering a rationale for its inverse effects on these subtypes. CGP36742 binding is similar to that of TPMPA, detailing a shared mechanism of action among phosphinic acid inhibitors. In contrast, GABOB binding is similar to that of GABA, but under equivalent conditions produces a mixture of primed and desensitized states; its density is compatible with both enantiomeric forms, likely representing the racemic mixture. Together, these results elucidate detailed interactions of a ρ-type GABA_A_ receptor with therapeutic drugs, offering mechanistic insights and a prospective basis for further drug development.

## Results

### Distinct states of ρ1 GABA_A_ receptors resolved with antagonist and agonist drugs

We implemented a modified human ρ1 GABA_A_ receptor construct (ρ1-EM) with truncated loops in the N-terminus and intracellular domain and an inserted fluorescent protein to facilitate expression while largely preserving wild-type function^27^. In *Xenopus laevis* oocytes expressing ρ1-EM, THIP and CGP36742 antagonized GABA activation (Fig.1a,b), consistent with previous studies of wild-type channels^28, 29^. GABOB functioned as an agonist, albeit at ∼10-fold higher concentrations than GABA (Fig.1c), consistent with its relatively weaker activity^30^.

**Figure 1.**
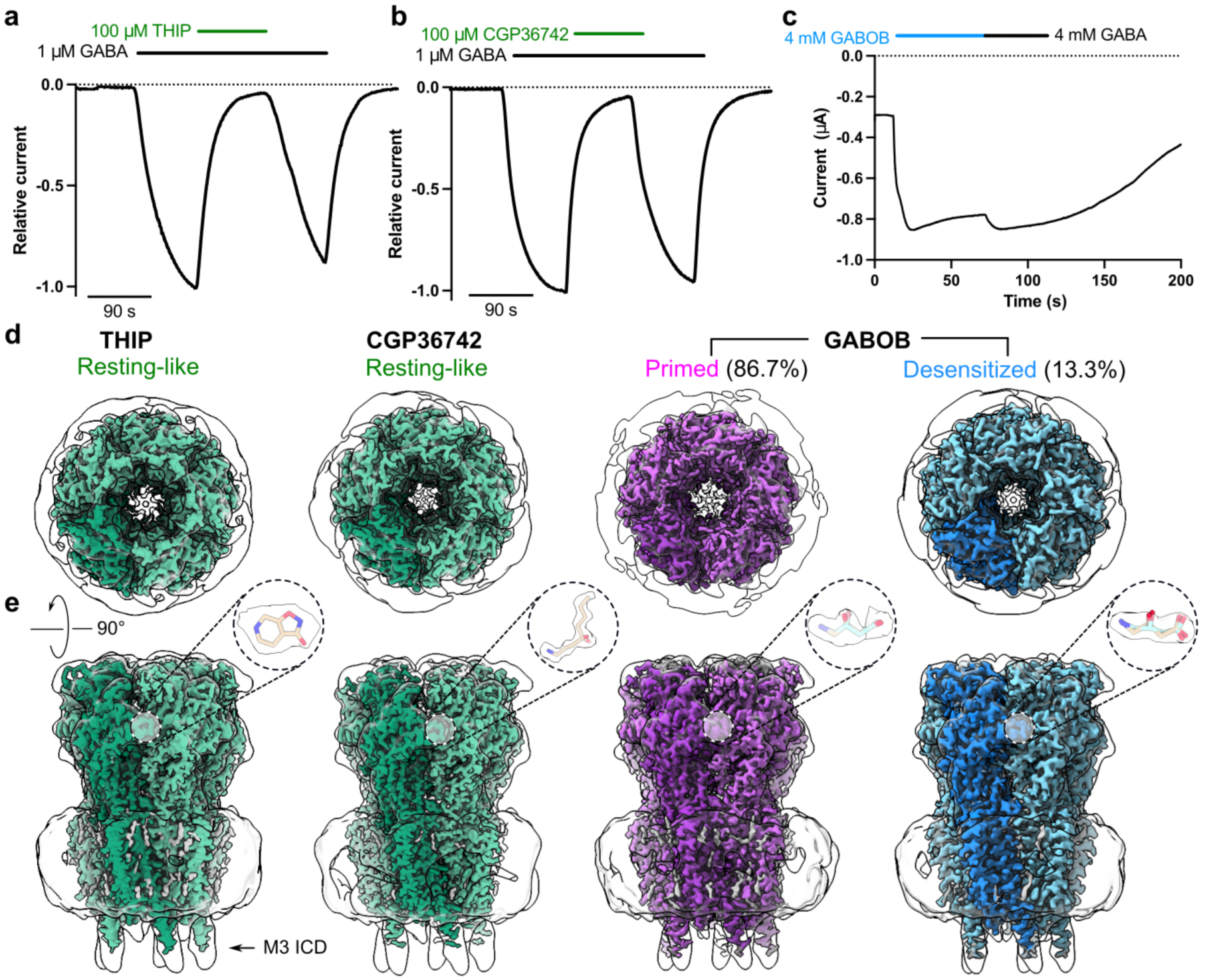
Functional and structural profiles of ρ1 with antagonist and agonist drugs. (a-c) Sample traces from two-electrode voltage-clamp electrophysiology recordings of ρ1-EM expressed in *Xenopus oocytes* exposed to THIP, CGP36742 or GABOB. (d) Cryo-EM maps of human ρ1-EM in complex with THIP (left), CGP36742 (middle) or GABOB (right), viewed from the extracellular side. Maps are colored by assigned functional states as similar to resting (green), primed (purple), or desensitized (blue). Low-pass filtered maps are shown in transparency to reveal lower-resolution features including the nanodisc and M3-helix intracellular-domain extension (M3 ICD). (e) Cryo-EM maps as in *d*, viewed from the membrane plane. Insets show the drugs with corresponding densities.

To gain insight into the binding and conformational changes associated with these drugs, we solved cryogenic electron microscopy (cryo-EM) structures of ρ1-EM in complex with THIP, CGP36742 and GABOB in saposin nanodiscs with polar brain lipids to resolutions 2.0-2.4 Å (Fig.1, Supplementary Fig.2, 3, 4, Table 1). We observed non-protein densities corresponding to the expected drugs in the five extracellular orthosteric ligand-binding sites of each structure; the corresponding maps enabled us to unambiguously build models for both the protein and drugs (Fig.1, Supplementary Fig.5).

**Table 1.**
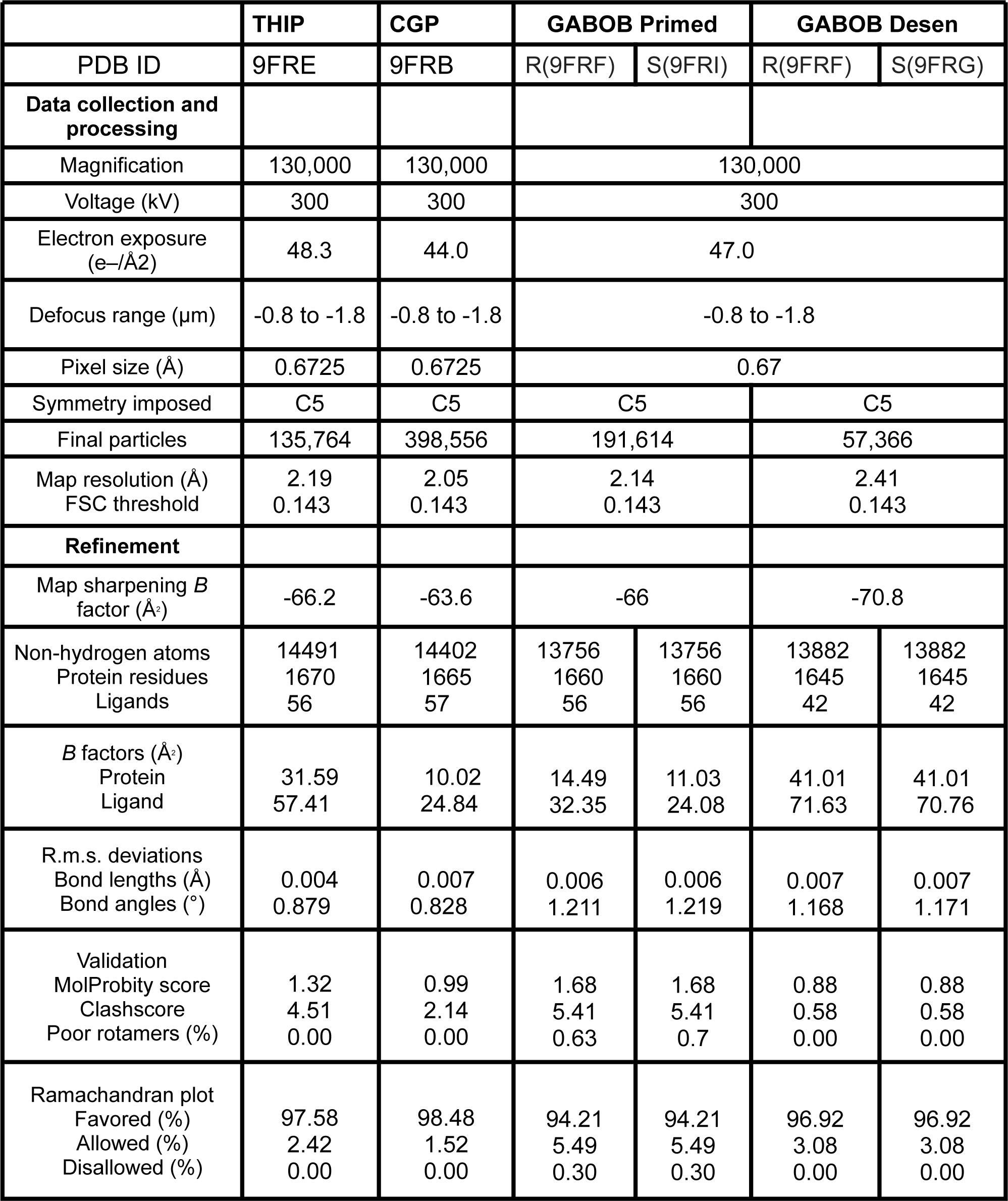
Cryo-EM data collection, refinement and validation statistics.

|  | THIP | CGP | GABOB Primed |  | GABOB Desen |  |
| --- | --- | --- | --- | --- | --- | --- |
| PDB ID | 9FRE | 9FRB | R(9FRF) | S(9FRI) | R(9FRF) | S(9FRG) |
| Data collection and processing |  |  |  |  |  |  |
| Magnification | 130,000 | 130,000 | 130,000 |  |  |  |
| Voltage (kV) | 300 | 300 | 300 |  |  |  |
| Electron exposure (e-/Å2) | 48.3 | 44.0 | 47.0 |  |  |  |
| Defocus range (µm) | -0.8 to -1.8 | -0.8 to -1.8 | -0.8 to -1.8 |  |  |  |
| Pixel size (Å) | 0.6725 | 0.6725 | 0.67 |  |  |  |
| Symmetry imposed | C5 | C5 | C5 |  | C5 |  |
| Final particles | 135,764 | 398,556 | 191,614 |  | 57,366 |  |
| Map resolution (Å) | 2.19 | 2.05 | 2.14 |  | 2.41 |  |
| FSC threshold | 0.143 | 0.143 | 0.143 |  | 0.143 |  |
| Refinement |  |  |  |  |  |  |
| Map sharpening *B* factor (Å²) | -66.2 | -63.6 | -66 |  | -70.8 |  |
| Non-hydrogen atoms | 14491 | 14402 | 13756 | 13756 | 13882 | 13882 |
| Protein residues | 1670 | 1665 | 1660 | 1660 | 1645 | 1645 |
| Ligands | 56 | 57 | 56 | 56 | 42 | 42 |
| *B* factors (Å²) |  |  |  |  |  |  |
| Protein | 31.59 | 10.02 | 14.49 | 11.03 | 41.01 | 41.01 |
| Ligand | 57.41 | 24.84 | 32.35 | 24.08 | 71.63 | 70.76 |
| R.m.s. deviations |  |  |  |  |  |  |
| Bond lengths (Å) | 0.004 | 0.007 | 0.006 | 0.006 | 0.007 | 0.007 |
| Bond angles (°) | 0.879 | 0.828 | 1.211 | 1.219 | 1.168 | 1.171 |
| Validation |  |  |  |  |  |  |
| MolProbity score | 1.32 | 0.99 | 1.68 | 1.68 | 0.88 | 0.88 |
| Clashscore | 4.51 | 2.14 | 5.41 | 5.41 | 0.58 | 0.58 |
| Poor rotamers (%) | 0.00 | 0.00 | 0.63 | 0.7 | 0.00 | 0.00 |
| Ramachandran plot |  |  |  |  |  |  |
| Favored (%) | 97.58 | 98.48 | 94.21 | 94.21 | 96.92 | 96.92 |
| Allowed (%) | 2.42 | 1.52 | 5.49 | 5.49 | 3.08 | 3.08 |
| Disallowed (%) | 0.00 | 0.00 | 0.30 | 0.30 | 0.00 | 0.00 |

Consistent with their functional roles as competitive inhibitors, THIP- and CGP36742-bound structures of ρ1-EM adopted the resting-like state, nearly identical to our previously reported apo structure^27^ (Table 2). In the presence of the agonist GABOB, one class (13% of resolved particles) corresponded to the desensitized state previously reported with GABA^27^ (Table 2). In another class (87% of resolved particles), despite the presence of GABOB in the orthosteric site, the ECD was only modestly activated, and the pore remained closed. This structure, corresponding neither to the resting nor desensitized states, aligned well with a previous so-called primed state determined in the presence of the negative modulator 17β-estradiol (E2)^31^ (Supplementary Fig.4e,f) (Table 2). Notably, E2 binds in a pocket located at the ECD-TMD interface, where it apparently disrupts allosteric transitions induced by GABA binding. In contrast, we observed no ligand density at the equivalent interface in the GABOB-bound structures, indicating a distinct mechanism of primed-state stabilization.

**Table 2.**
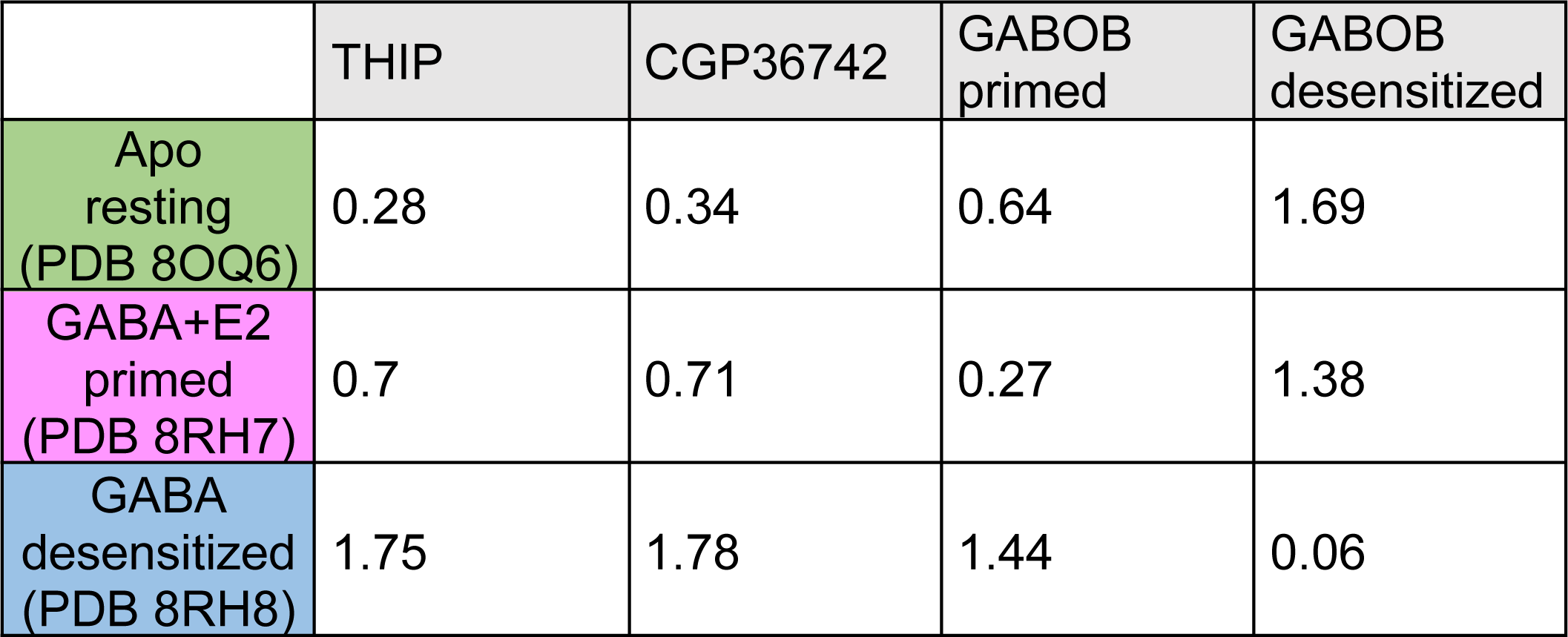
Cɑ RMSD (Å) between the ρ1-EM structures.

### Distinctive binding poses implicated in subtype-dependent effects of THIP

In our cryo-EM structure, THIP was bound in the orthosteric site with its two heterocycles perpendicular to loop C (Fig.2a). The pyridine ring faced the principal subunit, with its amino group buried in an aromatic cage involving residues Y219, Y262 and Y268. The isoxazole ring faced the complementary subunit, with its hydroxyl and amino groups making polar interactions with principal residues S264 and T265, as well as complementary residues R125 and S189 (Fig.2a,b, Supplementary Fig.6).

**Figure 2.**
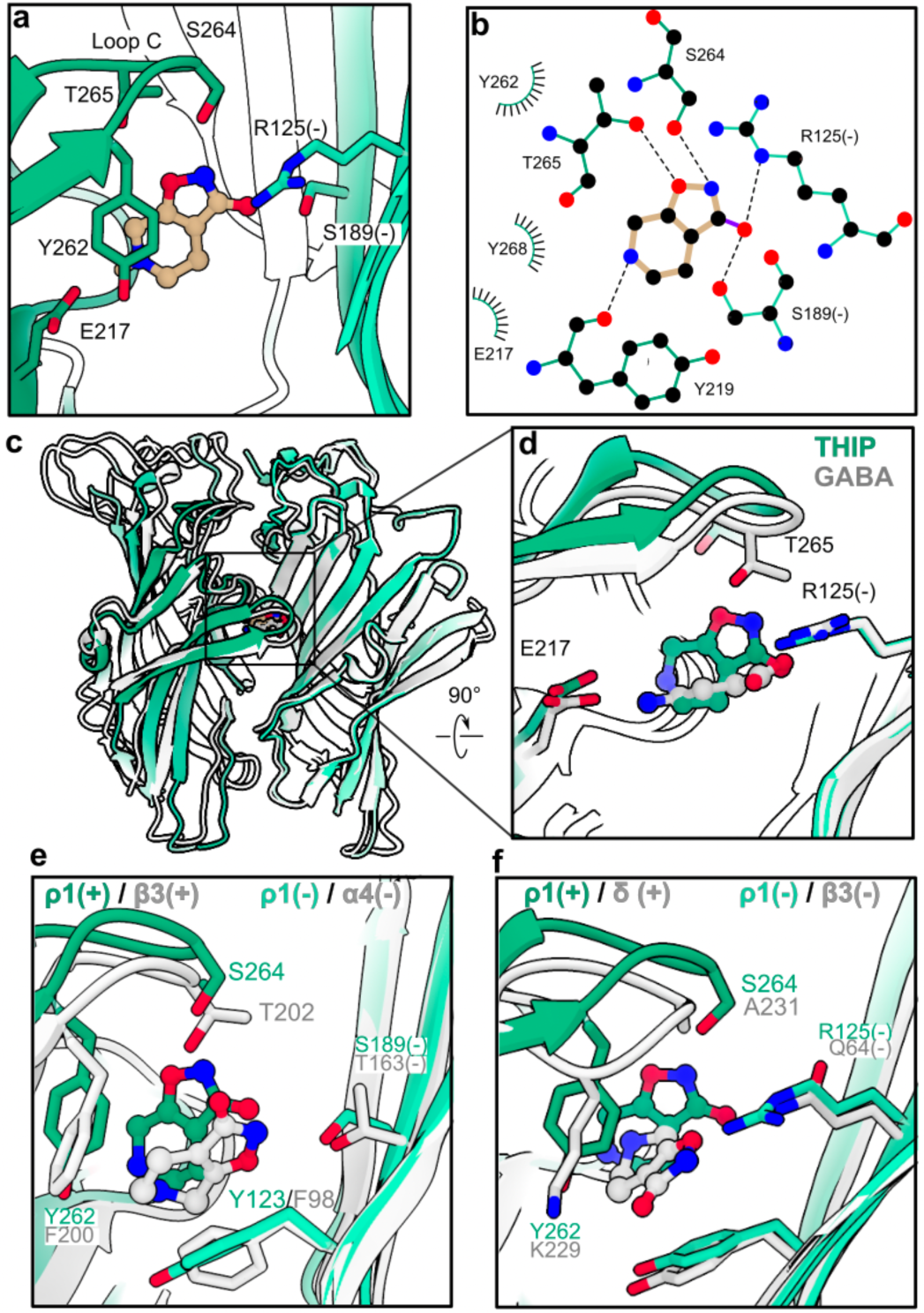
Distinctive binding pose of THIP in the ρ1 GABA_A_ receptor. (a) Zoom view of the THIP (tan) binding site in ρ1-EM (green). Residues interacting directly with THIP are shown as sticks and colored by heteroatom; residues from the complementary subunit face are denoted (-). (b) Schematic of ρ1-EM interactions with THIP, colored as in *a*. Hydrogen bonds are indicated as dashed lines, hydrophobic and aromatic interactions as lashes. (c-d) Superimposition of the ECD of THIP-bound (green) and GABA-bound (PDB ID 8OP9, gray) ρ1-EM structures. For clarity, only two subunits are shown, aligned on the complementary subunit. (e-f) Zoom views as in *d* of the superimposition of the THIP binding site in ρ1-EM (green) and the α4β3δ GABA_A_ receptor (PDB ID 7QND, gray), focusing on the β3-ɑ4 or δ-β3 interfaces, respectively. Structures are aligned on the ECD of the complementary subunit.

To visualize structural changes induced by antagonist versus agonist binding, we superimposed the resting-like THIP complex with our previous GABA-bound desensitized structure^27^ (Fig.2c,d). The amino and hydroxyl groups of THIP roughly aligned with those of GABA; however, the bulky heterocycles of THIP were relatively protruded toward loop C, obstructing its lockdown over the antagonist. Thus, similar to TPMPA^27^, THIP appears to sterically occlude local activating transitions in the ρ-type orthosteric site.

Whereas THIP is a competitive antagonist of ρ1^32^ (Fig.1a), it is a partial agonist of α1β3γ2 GABA_A_ receptors associated with the postsynapse, and a super-agonist of α4β3δ GABA_A_ receptors found extrasynaptically^33–35^. Consistent with this behavior, a recent cryo-EM structure of the α4β3δ subtype was reported with THIP in an apparent desensitized state, with ligands at both β3(+)-α4(-) and δ(+)-β3(-) interfaces^36^. At both these interface types, THIP adopts a pose distinct from that in the ρ1 resting state, with the heterocycles rotated nearly 90° to lie parallel to the plane of loop C (Fig.2e,f). This THIP pose in α4β3δ GABA_A_ receptors is compatible with more extensive lockdown of loop C than in ρ1, apparently enabling activation. We previously observed that loop C is extended by one residue in ρ1 relative to β GABA_A_-receptor subunits, and that truncating the amino acid at the tip of loop C (ΔS264) results in functional properties more similar to postsynaptic subtypes^27^. THIP effects were similarly decreased in the ΔS264 variant, but remained inhibitory (Supplementary Fig.7a), indicating that factors other than loop C length are involved in subtype-specific modulation. Although the structural basis for the distinct binding pose is not entirely clear, several bulky groups in the ρ1 orthosteric site are substituted with smaller sidechains in β3, including Y262 (β3-F200) on the principal subunit and R125 (β3-Q64) and S189 (β3-G127) on the complementary subunit (Supplementary Fig.6).

### Structural basis for receptor specificity among phosphinic acid inhibitors

In our cryo-EM structure, CGP36742 adopted an elongated pose, with its aminopropyl tail forming a salt bridge with E217 and cation-π interactions with the aromatic cage on the principal subunit face. The central phosphinic acid group was wedged between loop C on the principal subunit (making polar interactions with S264 and T265) and residue R125 on the complementary subunit, with the butyl tail extending toward the complementary β5 strand (Fig.3a,b). The complex was comparable to the apo and THIP structures (Table 2, Fig.3c), and nearly superimposable with our previous TPMPA structure (Fig.3d), including analogous interactions of the amino and phosphinic acid groups; the additional butyl tail only required an alternative rotamer of M177 in the β5 strand (Fig.3c,d). As in the case of TPMPA, the bulky, electronegative phosphinic acid group appeared to prevent lockdown of loop C over the antagonist, indicating a shared mechanism as well as binding pose among phosphinic acid inhibitors (Fig.3e,f).

**Figure 3.**
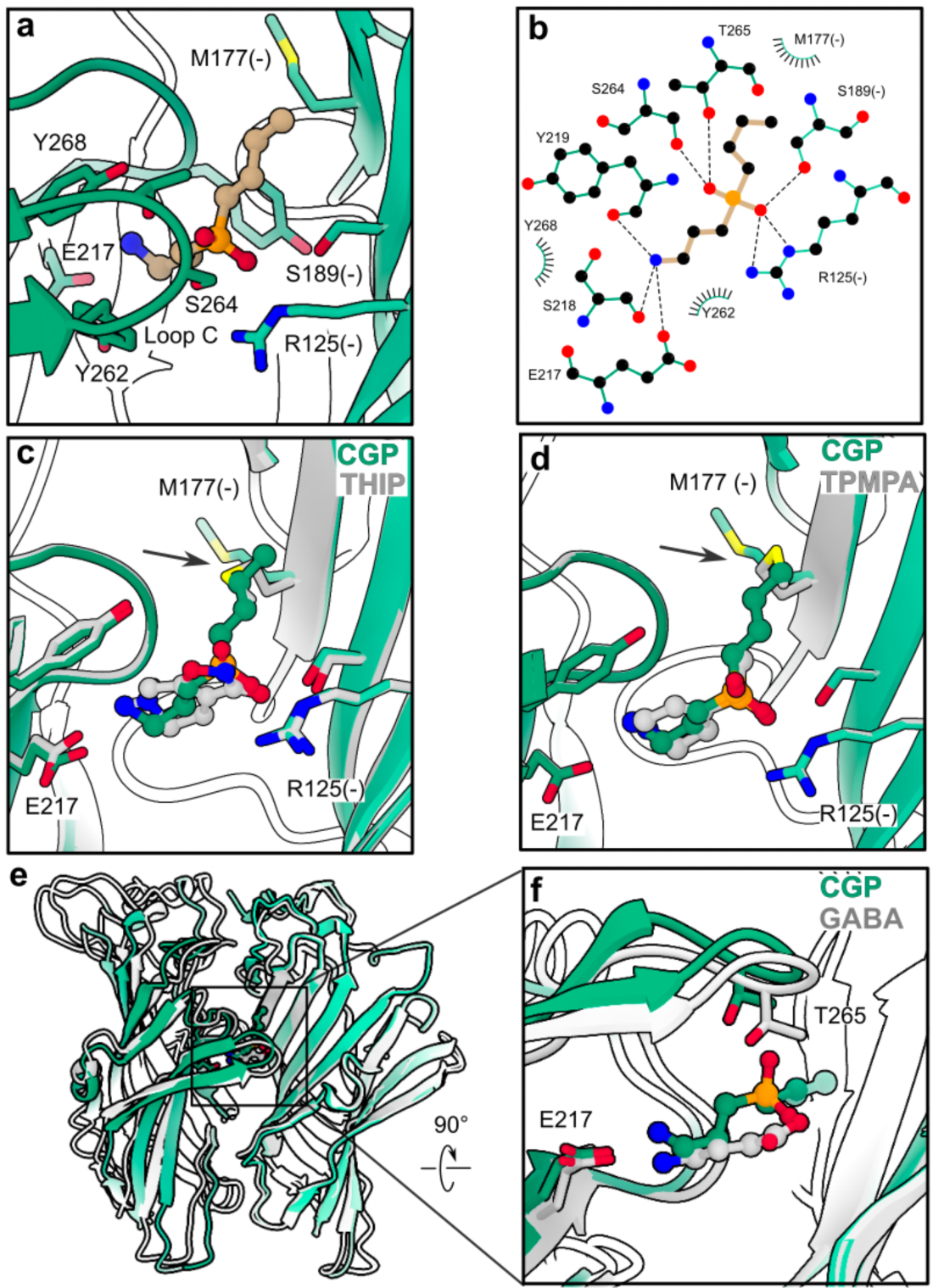
Common mechanism of ρ1 GABA_A_ receptor antagonism by phosphinic acid inhibitors. (a) Zoom view of CGP36742 (tan) binding site in ρ1-EM (green), colored and labeled as in *Fig.2a*. (b) Schematic of ρ1-EM interactions with CGP36742. Hydrogen bonds and other electrostatic interactions are indicated as dashed lines, hydrophobic and aromatic interactions as lashes. (c) Zoom view of superimposed structures of ρ1-EM with CGP36742 (green) and THIP (gray). (d) Zoom view of superimposed structures of ρ1-EM with CGP36742 (green) and TPMPA (PDB ID 8OQ7, gray). (e-f) Superimposition of the ECD of CGP36742-bound (green) and GABA-bound (PDB ID 8OP9, gray) ρ1-EM structures. For clarity, only two subunits are shown, aligned on the complementary subunit.

Given the potency of CGP36742 to inhibit GABA_B_ as well as ρ1 GABA_A_ receptors^29, 37^, we also compared its potential interactions with these structurally distinct targets. A structure of the related phosphinic acid inhibitor CGP35348 was previously reported with the ECD of a human GABA_B_ receptor, bound in the cleft between ligand-binding lobes (LB1 and LB2) of the GBR1 subunit^38^ (Fig.4a-c). The CGP36742 pose from our ρ1-EM structure could be aligned into this GABA_B_ receptor binding pocket on its equivalent aminopropyl and phosphinic acid moieties without clashing with any surrounding residues (Fig.4b). Interestingly, the amino group of CGP36742 was oriented roughly opposite to that of CGP35348 in its resolved site (Fig.4d), indicating the rotational flexibility of these compounds could support their polymodal activities.

**Figure 4.**
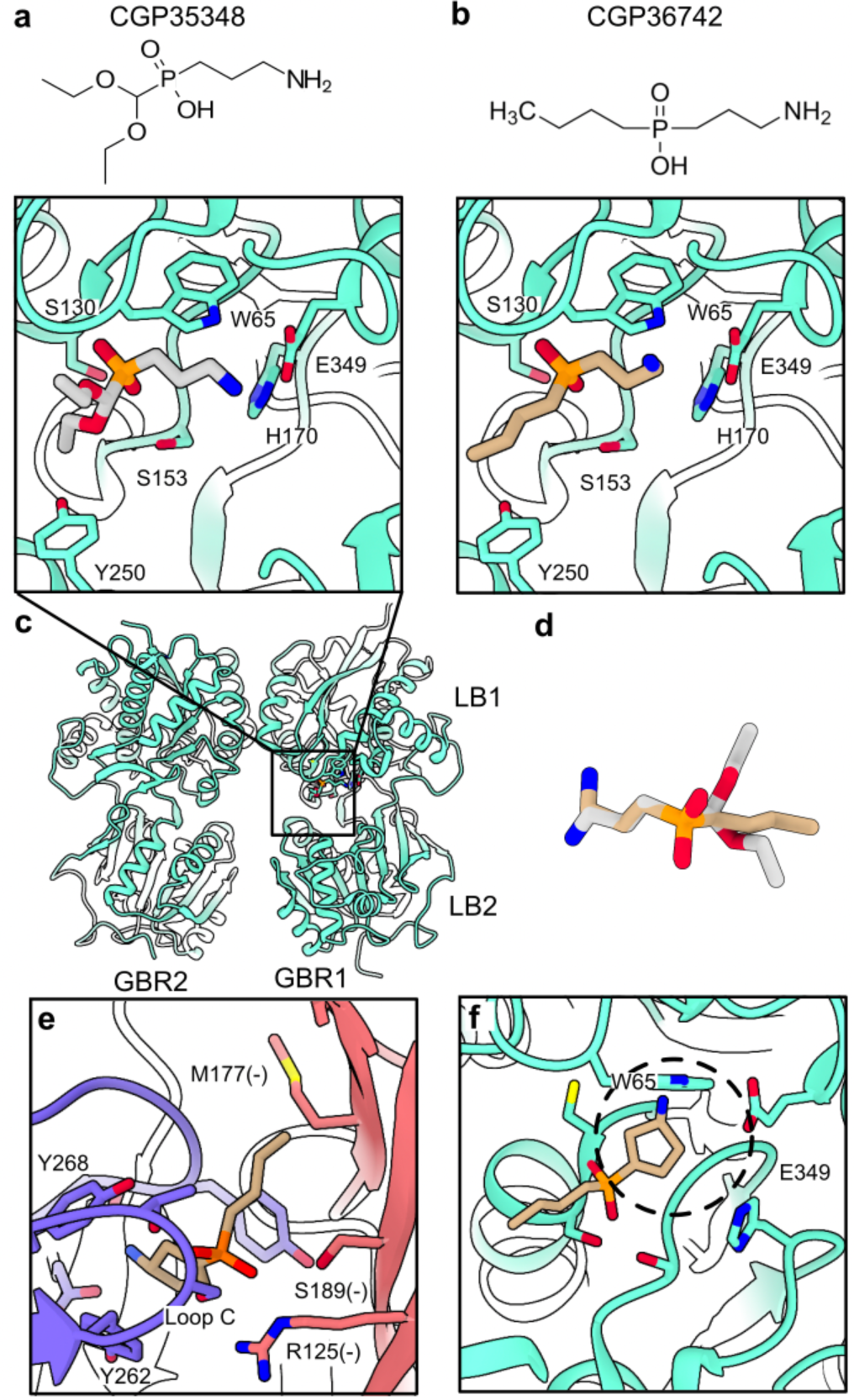
Comparative interactions of GABA_B_ and GABA_A_ receptors with phosphinic acid inhibitors. (a) Chemical structure of CGP35348 (above) and its binding site in a human GABA_B_ receptor (PDB ID 4MR8, green, below). The ligand (gray) and its direct residue contacts are colored by heteroatom. (b) Chemical structure of CGP36742 (above), and its superimposition into the equivalent GABA_B_-receptor site as in *a*. The ligand pose is adopted from the structure reported here with ρ1-EM, aligned on the shared aminopropyl and phosphinic acid moieties of CGP35348. (c) Overview of the ECD of the human GABA_B_ receptor as shown in (a). (d) Alignment of CGP36742 in ρ1-EM (tan) with CGP35348 in the GABA_B_ receptor (gray) as implemented in (b), colored by heteroatom. (e) Alignment of (S)-4-ACPBPA (tan) into the CGP36742 site of ρ1-EM. For perspective, principal and complementary subunits of ρ1-EM are colored purple and pink respectively; the ligand and interacting residues are colored by heteroatom. (f) Alignment of (S)-4-ACPBPA (tan) into the CGP35348 of the GABA_B_ receptor site (green) shown in (a). Dashed circle indicates prospective clash with residue W65.

The most potent and selective ρ1 antagonist identified thus far, (4-aminocyclopenten-1-yl)-butylphosphinic acid ((S)-4-ACPBPA), shares the phosphinic acid and butyl groups of CGP36742 but has a conformationally restricted aminocyclopentenyl group in place of the flexible aminopropyl tail^39, 40^ (Supplementary Fig.1). Aligning (S)-4-ACPBPA with CGP36742 in our ρ1-EM complex showed this ligand could be accommodated without modification (Fig.4e). Conversely, aligning this compound into the GABA_B_ receptor resulted in a clash of the amino group with residue W65, suggesting a molecular basis for receptor specificity (Fig.4f).

### Primed and desensitized states captured with the weak racemic agonist GABOB

As described above, our cryo-EM dataset collected with GABOB contained particles in both apparent primed and desensitized states. The primed state exhibited only a partial lockdown of loop C over the ligand, corresponding to a limited (1.8 Å) S264-Cα translation, and a subtle (1.2°) domain rotation relative to the resting state (Fig.5a). Although the contribution of this primed state to the receptor gating cycle remains unclear, the TMD was superimposable with that of the resting state, consistent with it representing a pre-active intermediate between resting and open. The presence of a substantial primed class with GABOB may reflect the relatively low affinity and slow kinetics of this agonist, despite the application of a supersaturating concentration (4 mM) for >30 min prior to grid freezing.

**Figure 5.**
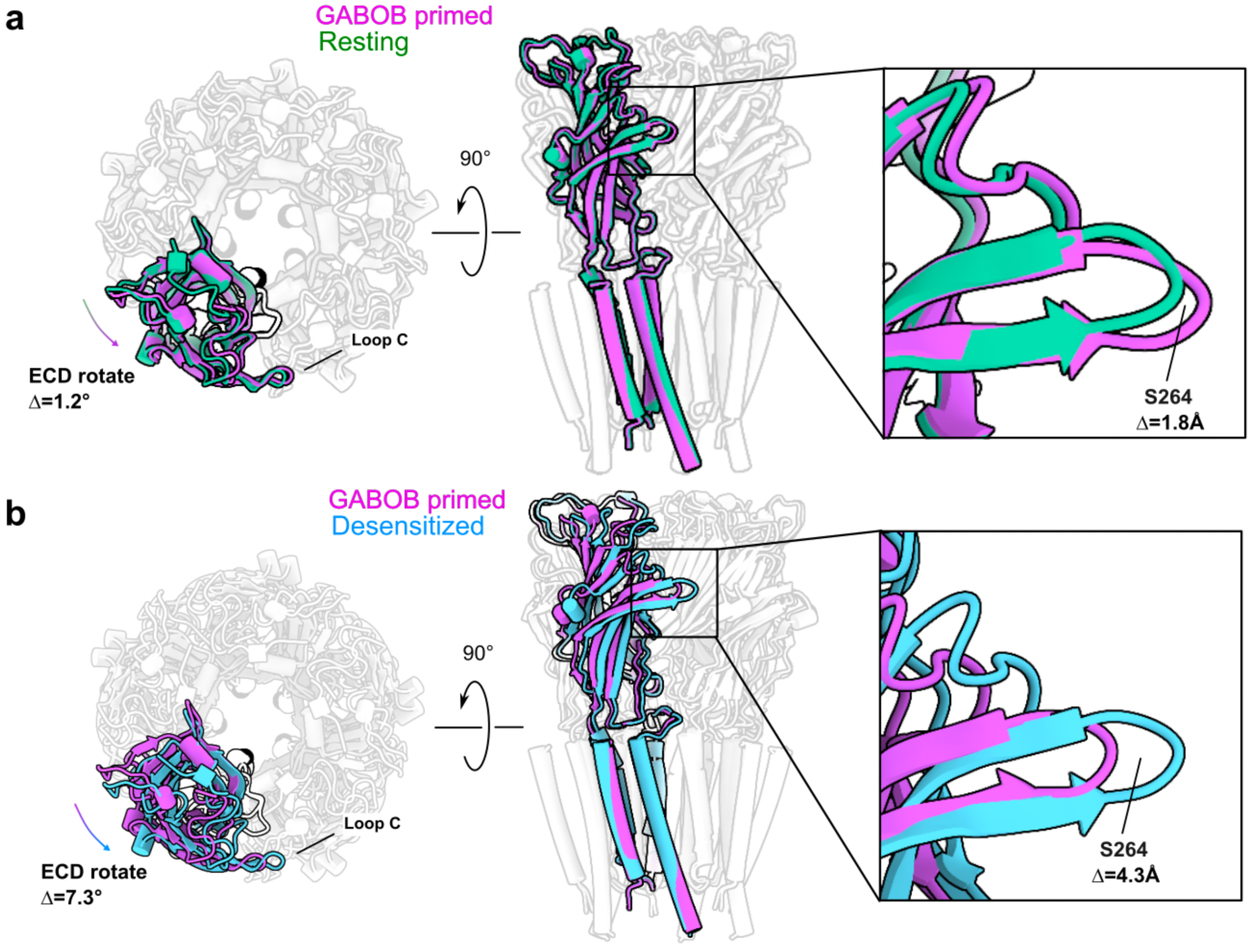
Primed and desensitized states captured with GABOB. (a) Superimposed structures of the GABOB-bound primed state (purple) with a previously reported resting state (green, PDB 8OQ6). Structures are aligned on the TMD; for clarity, all but one of the subunits are semi-transparent. (b) Superimposed structures of GABOB-bound primed state (purple) and desensitization state (blue). The structures are aligned with the TMD domain, and for clarity all but one of the subunits are transparent.

Consistent with our previous structure with GABA^27^, the desensitized state with GABOB exhibited further lockdown of loop C (4.3 Å S264-Cα translation) and rotation between the ECD and TMD (7.3°) relative to the primed state (Fig.5b). Interestingly, although all structures in this work contained ligands in the extracellular orthosteric site, local resolution in the ECD was relatively higher in the GABOB-desensitized state; in resting and primed structures, resolution was roughly similar between the domains (Supplementary Fig.4a-d), suggesting that channel activation is associated with both stabilization of the ECD and mobilization of the TMD.

Densities for GABOB were clearly resolved in both the primed and desensitized states (Fig.6a,b). Although the C3 carbon of GABOB constitutes a stereocenter, both enantiomers are ρ1 agonists, with modestly greater potency for (R)-GABOB^30^. As in medical practice, we used a racemic mixture in our experiments, such that our cryo-EM densities likely contained both forms; indeed, either (R)- or (S)-GABOB could be modeled into the ligand density in either structure (Fig.6a,b). The amino end was coordinated by an aromatic cage (Y219, Y262, Y268) as well as by E217 in the principal subunit, while the carboxylate end made electrostatic contacts with R125 and S189 in the complementary subunit (Fig.6c-f). In all cases, T265 at the tip of loop C could make a hydrogen bond with the C3 hydroxyl, consistent with the demonstrated effect of this residue on potency of both GABOB forms^41^. To substantiate binding capacity for both entantiomers, we performed all-atom molecular dynamics (MD) simulations of both the primed- and desensitized-state structures with both (R)- and (S)-GABOB. In quadruplicate ≥400-ns simulations of each system, both enantiomers remained within 2 Å median root-mean-square deviation (RMSD) relative to their starting poses in both structures (Fig.6g, Supplementary Fig.8), consistent with weak enantiomeric specificity.

**Figure 6.**
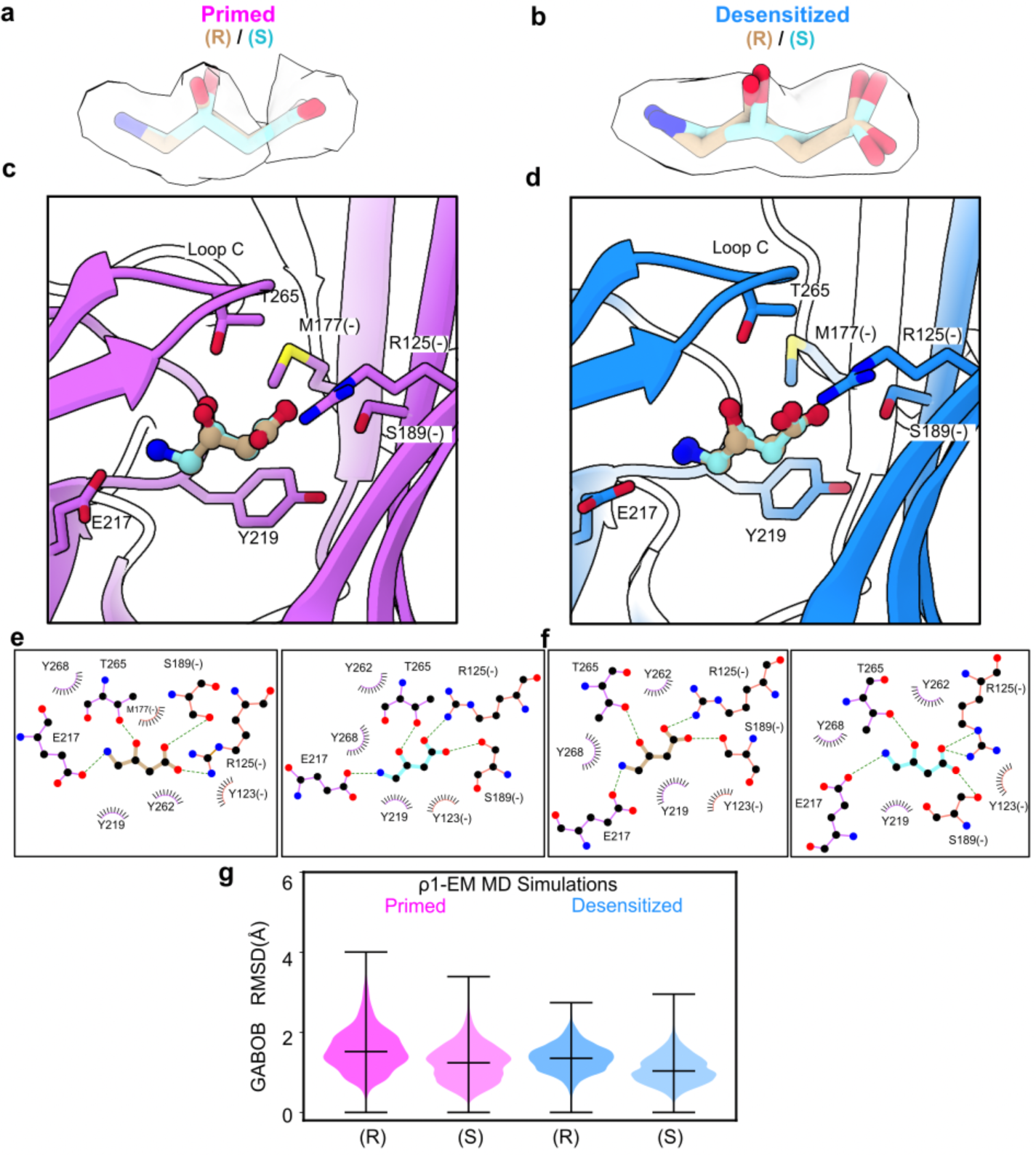
Comparable accommodation of GABOB enantiomers in the orthosteric site. (a) Modeling of (R)- (tan) and (S)-GABOB (cyan) into ligand density in the primed state of ρ1-EM (purple), colored by heteroatom. (b) Modeling of (R)- and (S)-GABOB into ligand density in the desensitized state of ρ1-EM (blue), otherwise colored as in *a*. (c) Zoom view of (R)- and (S)-GABOB in the primed state of ρ1-EM, colored as in *a*. (d) Zoom view of (R)- and (S)-GABOB in the desensitized state of ρ1-EM, colored as in *b*. (e) Schematics of (R)- (left, tan) and (S)-GABOB (right, cyan) interactions with the primed state of ρ1-EM. (f) Schematics of (R)- (left, tan) and (S)-GABOB (right, cyan) interactions with the desensitized state of ρ1-EM. In E-F, hydrogen bonds and other electrostatic interactions are indicated as dashed lines, hydrophobic and aromatic interactions as lashes. (g) Mobility of (R)- and (S)-GABOB in MD simulations of ρ1-EM in the primed (left, purple) and desensitized states (right, blue), as quantified by ligand RMSD (Å). Violin plots represent probability densities from 4 independent simulation replicates, sampled every 0.4 ns for the first 400 ns of each replicate (n = 4000), with markers indicating median and extrema.

## Discussion

Pharmaceutical targeting of ρ-type GABA_A_ receptors holds promise for drug development in treating visual, sleep, learning and memory disorders^13, 42^. Given the limited sensitivity of this subtype to classical GABA_A_ receptor modulators such as benzodiazepines, barbiturates and general anesthetics, the development of such agents will likely require a detailed understanding of ρ-specific mechanisms including binding, activation and inhibition. However, the specific mechanisms also means such drugs could limit interactions on neuronal receptors. The structural, functional and computational work presented here uncovers the binding modes of three drugs in the orthosteric site, highlighting among other things the critical role of loop C lockdown in channel gating.

Our structures highlight the critical role for loop-C lockdown in initiating ρ1 activation. Agonists such as GABA enable substantial lockdown of loop C, resulting in compaction of the orthosteric site and rotation of the ECD relative to the TMD (Fig.7a). Alongside this apparent activated-desensitized state, treatment with the weaker agonist GABOB promotes a subclass in a presumably intermediary primed state, similar to a structure previously reported with the inhibitor estradiol, and exhibiting only a modest lockdown of loop C relative to the apo form. It remains unclear whether the predominance of the primed state arises from partial occupancy of the relatively low-potency agonist GABOB, or its limited efficacy under cryo-EM conditions; or indeed, why a fully activated-open state of ρ1 remains inaccessible under cryo-EM conditions^27, 31^. On the other hand, antagonists such as THIP and phosphinic acids clearly obstruct loop-C lockdown altogether, resulting in an expanded orthosteric site superimposable with that of the apo structure (Fig.7b,c). Among phosphinic acid inhibitors, lockdown is obstructed by a consistent binding pose of CGP36742 and TPMPA, and likely of the subtype-specific agent (S)-4-ACPBPA. Conversely, cross-reactivity of CGP36742 with GABA_B_ receptor is attributable at least in part to rotational flexibility around the amino group.

**Figure 7.**
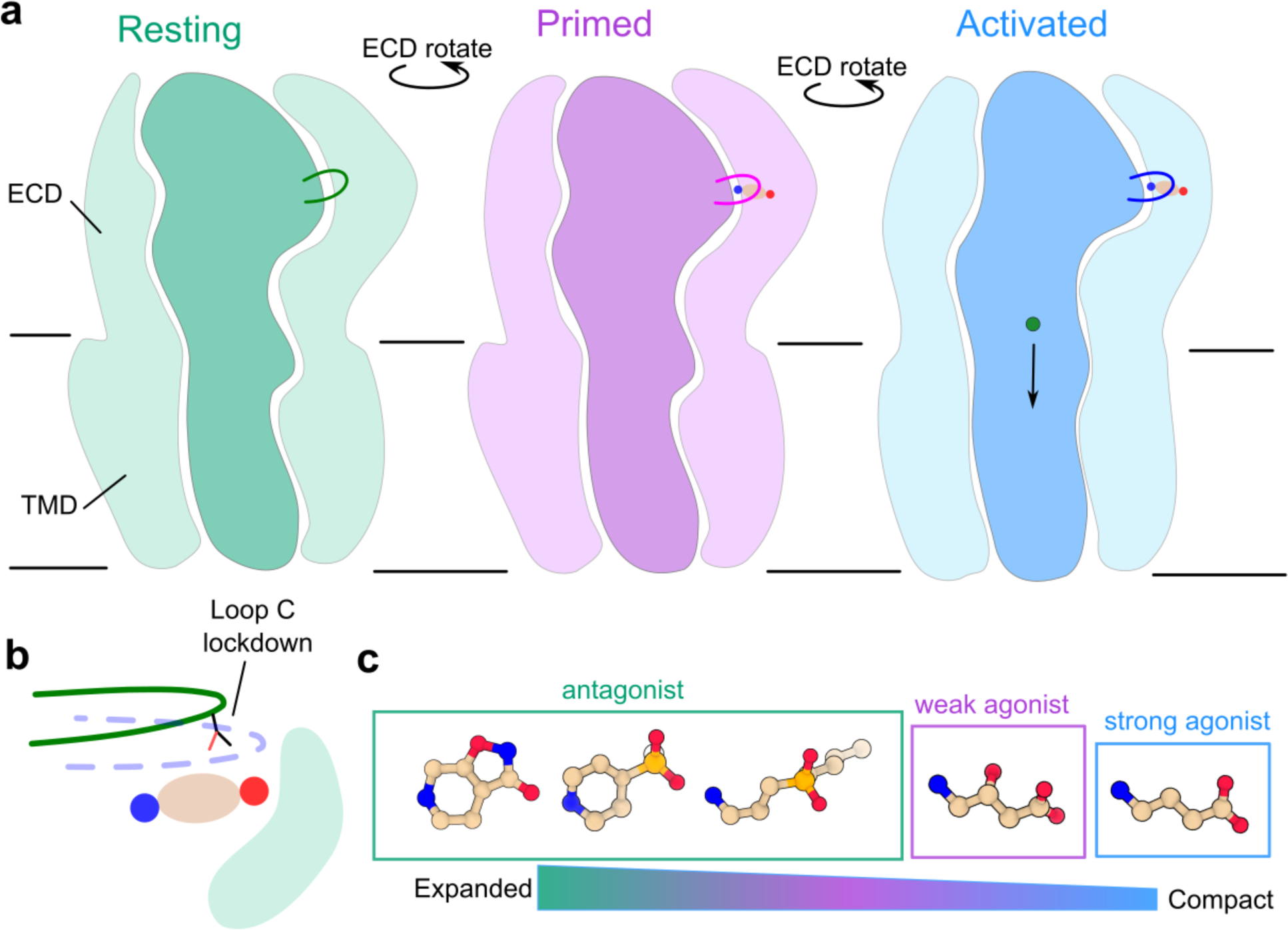
Proposed mechanisms of antagonist and agonist drugs. (a) Cartoons of ρ1 showing progressive ECD rotation between resting (green), primed (purple) and activated (open or desensitized, blue) states. Loop C is highlighted, and GABA and ions are shown in circles (carbon, tan; amine, blue; carboxylate, red; chloride, green). (b) Cartoon zoom view of the orthosteric binding site. GABA is shown in circles as in *a*. Green and blue lines depict lockdown of loop C. (c) Proposed dependence of orthosteric-site expansion/compaction on ligand identity, from relatively large antagonists to small agonists.

Subtype- and agent-specific interactions, e.g. with T265 on loop C, suggest avenues for future structure-based drug design. For instance, we observed a distinctive binding pose for THIP in ρ1-EM compared to a previously reported complex with an α4β3δ GABA_A_ receptor^36^, which may underlie its opposing effects in these subtypes. Interestingly, the THIP derivative aza-THIP is a more selective ρ-type antagonist, with activity comparable to THIP at ρ1 but negligible at heteromeric GABA_A_ receptors^28^. The two molecules are identical except at one heavy atom in the 5-membered ring, substituting a pyrazole in aza-THIP for the isoxazolo group in THIP (Supplementary Fig.1a). In structures with THIP, the substituted oxygen atom appears to accept a hydrogen bond from principal-subunit loop C (T265) in ρ1-EM, but from the complementary subunit (ɑ4-T163 or β3-Q64) in the α4β3δ GABA_A_ receptor. The differential environments for this substituted atom in particular may account for discriminating activity between these two receptors. A hydrogen bond with loop-C T265 also appears to underlie potency of both enantiomeric forms of GABOB in ρ1, and may contribute to the stabilization of a primed state in the presence of this weaker agonist relative to GABA. Taken together, this work details receptor-specific binding interactions of both antagonists and agonists in the orthosteric site, offering potential insights into differential pharmacology across multiple receptor subtypes in the GABAergic system.

## Methods

### Protein expression and purification

The expression and purification of ρ1-EM was following our earlier published methods^27^. Briefly, cell pellets from 2L of Expi293F cells infected with baculovirus were resuspended in the buffer (40 mM HEPES pH 7.5, 300 mM NaCl, with cOmplete protease inhibitor tablets (Roche)) and sonicated to break cell membranes. The membrane was pelleted by ultracentrifugation then resuspended and solubilized by resuspension buffer with 2% lauryl maltose neopentyl glycol (LMNG), 0.2% cholesteryl hemisuccinate (CHS) for 3 h in the cold room. The solubilization mixture was ultracentrifuged and the supernatant was applied to 4 mL Strep-Tactin XT Superflow resin (IBA) and incubated for 90 min. Resin was washed with wash buffer (20 mM HEPES pH 7.5, 300 mM NaCl, 0.005% LMNG, 0.0005% CHS) then protein was eluted with elution buffer (wash buffer with 10 mM d-Desthiobiotin (Sigma)). The product was further purified by size exclusion chromatography on a Superose 6 column (Cytiva) with flow buffer (20 mM HEPES pH 7.5, 100 mM NaCl, 0.005% LMNG, 0.0005% CHS). Peak fractions were pooled for nanodisc reconstitution.

### Nanodisc reconstitution

The plasmid for SapA expression was a gift from Salipro Biotech AB. Purification of SapA followed the published protocols^43^. For the reconstitution of nanodisc, ρ1-EM, SapA and polar brain lipid (Avanti) were mixed as molar ratio 1:15:150, then incubated on ice for 1 h. Bio-Beads SM-2 resin (Bio-Rad) was added into the mixture then gently rotated overnight at 4°C. On the next day, the supernatant was collected and further purified by gel-filtration chromatography on a Superose 6 column (Cytiva) with buffer containing 20mM HEPES pH 7.5, 100mM NaCl. Peak fractions were pooled and concentrated to ∼5 mg/mL.

### Cryo-EM grid preparation and data collection

The nanodisc sample was mixed with the compound stock solutions with volume ratio 9:1. The stock solutions were (5 mM THIP, 20mM fluorinated foscholine 8 (FFC-8)), (20 mM CGP36742, 20mM FFC-8), (40 mM GABOB, 20mM FFC-8). The mixtures were incubated on ice for more than 30 minutes before freezing grids. For each grid, 3 μL of the mixture was applied to a glow-discharged grid (R1.2/1.3 300 mesh Au grid, Quantifoil), blotted for 2 s with force 0 and plunged into liquid ethane using a Vitrobot Mark IV (Thermo Fisher Scientific). Cryo-EM data were collected on a 300kV Titan Krios (Thermo Fisher Scientific) electron microscope with a K3 Summit detector (Gatan) with magnification 130k corresponding to 0.6725 or 0.6645 Å/px using the software EPU 3.5.0 (Thermo Fisher Scientific). The total dose was ∼46 e^-^/Å^2^ and defocus range was −0.8 to −1.8 μm.

### Cryo-EM data processing

Dose-fractionated images in super-resolution mode were internally gain-normalized and binned by 2 in EPU during data collection. Cryo-EM data processing was first done in Relion 3.1.4^44^, including Motion correction, contrast transfer functions (CTF) estimation with CTFFIND 4.1^45^, automatic particle picking with topaz 0.2.5^46^, particle extraction, 2D classification, 3D classification, 3D refinement, CTF refinement and polishing. Briefly, two rounds of 2D classification were done to remove junk particles, 3D classification (3 classes) was used to analyze the structural heterogeneity. Particles from classes with protein features were centered and re-extracted, and were used for the 3D refinement with symmetry C5. Several rounds of CtfRefine and one round of polishing were executed to improve the resolution. The shiny particles were imported into CryoSPARC v4.2.1 for further processing^47^, including 3D classification with the PCA mode, and Non-Uniform Refinement^48^.

### Model building and refinement

Model building was started with rigid body fitting of the previously published resting (PDB ID 8OQ6), primed (PDB ID 8RH7) or desensitized (PDB ID 8RH8) state structure into the density. The models were manually checked and adjusted in Coot 0.9.5^49^, and chemicals, waters and lipids were also manually added. The resulting model was further optimized using real-space refinement in PHENIX 1.18.2^50^ and validated by MolProbity^51^. Crystallographic information files (cif) for ligands were generated from isomeric SMILES strings using Grade2^52^.

### Structural analysis

Pore radius profiles were calculated using CHAP 0.9.1^53^. Structure figures were prepared using UCSF ChimeraX 1.3^54^. ECD rotation was calculated as the dihedral angle between a) the Cα COM of the ECD (residues 97-280) of one subunit, b) the equivalent ECD residues of all subunits, c) the TMD (residues 281-479) including all subunits, and d) the equivalent TMD residues of one subunit.

### Expression in oocytes and electrophysiology

mRNA encoding the ρ1-EM GABA_A_ receptor was produced by in-vitro transcription using the mMessage mMachine T7 Ultra transcription kit (Ambion) according to the manufacturer protocol. *Xenopus laevis* oocytes (Ecocyte Bioscience) were injected with 30-50 ng mRNA and incubated 4-8 days at 13°C in post-injection solution (10 mM HEPES pH 8.5, 88 mM NaCl, 2.4 mM NaHCO3, 1 mM KCl, 0.91 mM CaCl2, 0.82 mM MgSO4, 0.33 mM Ca(NO3)2, 2 mM sodium pyruvate, 0.5 mM theophylline, 0.1 mM gentamicin, 17 mM streptomycin, 10,000 u/L penicillin) prior to two-electrode voltage clamp measurements.

For recordings, glass electrodes were pulled and filled with 3 M KCl to give a resistance of 0.5-1.5 MΩ and used to clamp the membrane potential of injected oocytes at −60 mV with an OC-725C voltage clamp (Warner Instruments). Oocytes were maintained under continuous perfusion with Ringer’s solution (123 mM NaCl, 10 mM HEPES, 2 mM KCl, 2 mM MgSO4, 2 mM CaCl2, pH 7.5) at a flow rate around 1.5 mL/min. Buffer exchange was accomplished by manually switching the inlet of the perfusion system to the appropriate buffer. For assessing GABOB efficacy, a gravity-fed perfusion system was used as previously described to improve kinetics of solution exchange. Currents were digitized at a sampling rate of 2 kHz and lowpass filtered at 10 Hz with an Axon CNS 1440A Digidata system controlled by pCLAMP 10 (Molecular Devices).

### Molecular dynamics simulation

Atomic coordinates of the ρ1-GABOB complex with GABOB built as either of two enantiomers were used as starting models for MD simulations. Each subunit was split into two chains for simulation, due to unresolved residues in the M3-M4 loop in the experimental structure. The simulation systems were set up in CHARMM-GUI^55^. The protein was embedded into a bilayer mimicking brain-lipid composition with the top leaflet containing 155 POPC, 24 POPE, and 38 cholesterol molecules and the bottom leaflet containing 65 POPC, 115 POPE, 26 POPS, and 32 cholesterol molecules. The protein-lipid complex was subsequently solvated with TIP3P water and 150 mM NaCl. The CHARMM36m forcefield^56^ was used to describe the protein. Parameters for (R)- and (S)-GABOB were generated using CGenFF^57^ in CHARMM-GUI. Cation-π specific NBFIX parameters were used to maintain appropriate ligand-protein interactions in the aromatic cage in the orthosteric binding site^58^.

Simulations were performed using GROMACS 2022.5^59^ at 300 K and 1 bar using the velocity-rescaling thermostat^60^ and Parrinello–Rahman barostat^61^. The LINCS algorithm was used to constrain the length of all bonds involving hydrogens^62^, and the particle mesh Ewald method^63^ was used to calculate long-range electrostatic interactions. The systems were energy minimized and then equilibrated for 20 ns, with the position restraints on the protein and neurosteroids gradually released. Four replicates, each > 400 ns, were simulated for each system as final unrestrained production runs.

Before analysis, MD simulation trajectories were aligned on the Cα atoms of the ECD using MDAnalysis^64^. Root-mean-square deviations (RMSD) of the non-hydrogen atoms of GABOB were calculated in VMD^65^, and data from the first 400 ns were combined and plotted as violin plots using Matplotlib^65^.

## Data Availability

The cryo-EM maps and corresponding atomic coordinates have been deposited in the Electron Microscopy Data Bank (EMDB) and the Protein Data Bank (PDB) with THIP (EMD-50712, PDB-9FRE), CGP36742 (EMD-50710, PDB-9FRB), (R)-GABOB in the primed state (EMD-50714, PDB-9FRH), (S)-GABOB in the primed state (EMD-50714, PDB-9FRI), (R)-GABOB in the desensitized state (EMD-50713, PDB-9FRF), and (S)-GABOB in the desensitized state (EMD-50713, PDB-9FRG). MD simulation trajectories and parameter files are available on Zenodo (https://zenodo.org/records/11549592).

## Acknowledgements

We thank members of Molecular Biophysics Stockholm for feedback on the project and manuscript, and staff at the Swedish National Cryo-EM Facility for data collection and support. The data was collected at the Cryo-EM Swedish National Facility funded by the Knut and Alice Wallenberg, Family Erling Persson and Kempe Foundations, SciLifeLab and Stockholm University. MD simulations were performed using the computing facilities of the Swedish National Infrastructure for Computing (SNIC 2022/3-40), and supported by BioExcel (EuroHPC grant no. 101093290). JC was supported by an EMBO Postdoctoral Fellowship, CF by grant FV-5.1.2-0523-19 from Stockholm University, and RJH, and EL by grants from the Knut and Alice Wallenberg Foundation (2023.0254), the Swedish Research Council (2019-02433, 2021-05806) and the Swedish e-Science Research Center.

## Author contributions

C.F. and J.C. performed the biochemistry, cryo-EM sample preparation and data processing. C.F. performed model building, refinement, structural analysis and MD simulations. J.C. performed electrophysiology. R.J.H. and E.L. supervised the project. All authors contributed to the manuscript writing and revision.

## References

1. Chebib, M. & Johnston, G. A. GABA-Activated ligand gated ion channels: medicinal chemistry and molecular biology. J. Med. Chem. 43, 1427–1447 (2000).

2. Lynagh, T. & Pless, S. A. Principles of agonist recognition in Cys-loop receptors. Front. Physiol. 5, 160 (2014).

3. Howard, R. J. Elephants in the Dark: Insights and Incongruities in Pentameric Ligand-gated Ion Channel Models. J. Mol. Biol. 433, 167128 (2021).

4. Bettler, B., Kaupmann, K., Mosbacher, J. & Gassmann, M. Molecular structure and physiological functions of GABA(B) receptors. Physiol. Rev. 84, 835–867 (2004).

5. Shimada, S., Cutting, G. & Uhl, G. R. gamma-Aminobutyric acid A or C receptor? gamma-Aminobutyric acid rho 1 receptor RNA induces bicuculline-, barbiturate-, and benzodiazepine-insensitive gamma-aminobutyric acid responses in Xenopus oocytes. Mol. Pharmacol. 41, 683–687 (1992).

6. Drew, C. A., Johnston, G. A. R. & Weatherby, R. P. Bicuculline-insensitive GABA receptors: Studies on the binding of (−)-baclofen to rat cerebellar membranes. Neurosci. Lett. 52, 317–321 (1984).

7. Johnston, G. Medicinal Chemistry and Molecular Pharmacology of GABA-C Receptors. Curr. Top. Med. Chem. 2, 903–913 (2002).

8. Rosas-Arellano, A., Estrada-Mondragón, A., Martínez-Torres, A. & Reyes-Haro, D. GABAA-ρ Receptors in the CNS: Their Functional, Pharmacological, and Structural Properties in Neurons and Astroglia. Neuroglia 4, 239–252 (2023).

9. Dong, C., Picaud, S. & Werblin, F. GABA transporters and GABAC-like receptors on catfish cone- but not rod-driven horizontal cells. J. Neurosci. 14, 2648 (1994).

10. Varman, D. R., Soria-Ortíz, M. B., Martínez-Torres, A. & Reyes-Haro, D. GABAρ3 expression in lobule X of the cerebellum is reduced in the valproate model of autism. Neurosci. Lett. 687, 158–163 (2018).

11. Zheng, W. et al. Function of gamma-aminobutyric acid receptor/channel rho 1 subunits in spinal cord. J. Biol. Chem. 278, 48321–48329 (2003).

12. Arnaud, C., Gauthier, P. & Gottesmann, C. Study of a GABAc receptor antagonist on sleep-waking behavior in rats. Psychopharmacology (Berl.) 154, 415–419 (2001).

13. Chebib, M. et al. Novel, potent, and selective GABAC antagonists inhibit myopia development and facilitate learning and memory. J. Pharmacol. Exp. Ther. 328, 448–457 (2009).

14. Cunha, C., Monfils, M.-H. & Ledoux, J. E. GABA(C) Receptors in the Lateral Amygdala: A Possible Novel Target for the Treatment of Fear and Anxiety Disorders? Front. Behav. Neurosci. 4, 6 (2010).

15. McCarson, K. E. & Enna, S. J. GABA pharmacology: the search for analgesics. Neurochem. Res. 39, 1948–1963 (2014).

16. Krogsgaard-Larsen, P., Frølund, B., Liljefors, T. & Ebert, B. GABA(A) agonists and partial agonists: THIP (Gaboxadol) as a non-opioid analgesic and a novel type of hypnotic. Biochem. Pharmacol. 68, 1573–1580 (2004).

17. Wafford, K. A. & Ebert, B. Gaboxadol--a new awakening in sleep. Curr. Opin. Pharmacol. 6, 30–36 (2006).

18. Budimirovic, D. B. et al. Gaboxadol in Fragile X Syndrome: A 12-Week Randomized, Double-Blind, Parallel-Group, Phase 2a Study. Front. Pharmacol. 12, 757825 (2021).

19. Keary, C. et al. Gaboxadol in angelman syndrome: A double-blind, parallel-group, randomized placebo-controlled phase 3 study. Eur. J. Paediatr. Neurol. EJPN Off. J. Eur. Paediatr. Neurol. Soc. 47, 6–12 (2023).

20. Froestl, W. et al. SGS742: the first GABA(B) receptor antagonist in clinical trials. Biochem. Pharmacol. 68, 1479–1487 (2004).

21. Chebib, M., Vandenberg, R. J., Froestl, W. & Johnston, G. A. Unsaturated phosphinic analogues of gamma-aminobutyric acid as GABA(C) receptor antagonists. Eur. J. Pharmacol. 329, 223–229 (1997).

22. Froestl, W. et al. Phosphinic acid analogues of GABA. 2. Selective, orally active GABAB antagonists. J. Med. Chem. 38, 3313–3331 (1995).

23. Mondadori, C., Möbius, H. J. & Borkowski, J. The GABAB receptor antagonist CGP 36,742 and the nootropic oxiracetam facilitate the formation of long-term memory. Behav. Brain Res. 77, 223–225 (1996).

24. Genkova-Papazova, M. G., Petkova, B., Shishkova, N. & Lazarova-Bakarova, M. The GABA-B antagonist CGP 36742 prevent PTZ-kindling-provoked amnesia in rats. Eur. Neuropsychopharmacol. J. Eur. Coll. Neuropsychopharmacol. 10, 273–278 (2000).

25. Johnston, G. A. GABAA receptor pharmacology. Pharmacol. Ther. 69, 173–198 (1996).

26. Nakao, J., Hasegawa, T., Hashimoto, H., Noto, T. & Nakajima, T. Formation of GABOB from 2-hydroxyputrescine and its anticonvulsant effect. Pharmacol. Biochem. Behav. 40, 359–366 (1991).

27. Cowgill, J. et al. Structure and dynamics of differential ligand binding in the human ρ-type GABAA receptor. Neuron 111, 3450–3464.e5 (2023).

28. Krehan, D. et al. Aza-THIP and related analogues of THIP as GABA C antagonists. Bioorg. Med. Chem. 11, 4891–4896 (2003).

29. Ng, C. K. L. et al. Medicinal chemistry of ρ GABAC receptors. Future Med. Chem. 3, 197–209 (2011).

30. Hinton, T., Chebib, M. & Johnston, G. A. R. Enantioselective actions of 4-amino-3-hydroxybutanoic acid and (3-amino-2-hydroxypropyl)methylphosphinic acid at recombinant GABA(C) receptors. Bioorg. Med. Chem. Lett. 18, 402–404 (2008).

31. Woodward, R. M., Polenzani, L. & Miledi, R. Characterization of bicuculline/baclofen-insensitive (rho-like) gamma-aminobutyric acid receptors expressed in Xenopus oocytes. II. Pharmacology of gamma-aminobutyric acidA and gamma-aminobutyric acidB receptor agonists and antagonists. Mol. Pharmacol. 43, 609–625 (1993).

32. Stórustovu, S. I. & Ebert, B. Pharmacological characterization of agonists at delta-containing GABAA receptors: Functional selectivity for extrasynaptic receptors is dependent on the absence of gamma2. J. Pharmacol. Exp. Ther. 316, 1351–1359 (2006).

33. Brown, N., Kerby, J., Bonnert, T. P., Whiting, P. J. & Wafford, K. A. Pharmacological characterization of a novel cell line expressing human alpha(4)beta(3)delta GABA(A) receptors. Br. J. Pharmacol. 136, 965–974 (2002).

34. Mortensen, M., Ebert, B., Wafford, K. & Smart, T. G. Distinct activities of GABA agonists at synaptic- and extrasynaptic-type GABAA receptors. J. Physiol. 588, 1251–1268 (2010).

35. Sente, A. et al. Differential assembly diversifies GABA(A) receptor structures and signalling. Nature 604, 190–194 (2022).

36. Nowak, G. et al. Antidepressant-like activity of CGP 36742 and CGP 51176, selective GABAB receptor antagonists, in rodents. Br. J. Pharmacol. 149, 581–590 (2006).

37. Geng, Y., Bush, M., Mosyak, L., Wang, F. & Fan, Q. R. Structural mechanism of ligand activation in human GABA(B) receptor. Nature 504, 254–259 (2013).

38. Gavande, N. et al. Novel Cyclic Phosphinic Acids as GABAC ρ Receptor Antagonists: Design, Synthesis, and Pharmacology. ACS Med. Chem. Lett. 2, 11–16 (2011).

39. Kumar, R. J. et al. Novel gamma-aminobutyric acid rho1 receptor antagonists; synthesis, pharmacological activity and structure-activity relationships. J. Med. Chem. 51, 3825–3840 (2008).

40. Yamamoto, I. et al. Differentiating enantioselective actions of GABOB: a possible role for threonine 244 in the binding site of GABA(C) ρ(1) receptors. ACS Chem. Neurosci. 3, 665–673 (2012).

41. Johnston, G., Chebib, M., Hanrahan, J. & Mewett, K. GABAC Receptors as Drug Targets. Curr. Drug Target -CNS Neurol. Disord. 2, 260–268 (2003).

42. Lyons, J. A., Bøggild, A., Nissen, P. & Frauenfeld, J. Saposin-Lipoprotein Scaffolds for Structure Determination of Membrane Transporters. Methods Enzymol. 594, 85–99 (2017).

43. Zivanov, J. et al. New tools for automated high-resolution cryo-EM structure determination in RELION-3. eLife 7, e42166 (2018).

44. Grant, T., Rohou, A. & Grigorieff, N. cisTEM, user-friendly software for single-particle image processing. eLife 7, (2018).

45. Bepler, T. et al. Positive-unlabeled convolutional neural networks for particle picking in cryo-electron micrographs. Nat. Methods 16, 1153–1160 (2019).

46. Punjani, A., Rubinstein, J. L., Fleet, D. J. & Brubaker, M. A. cryoSPARC: algorithms for rapid unsupervised cryo-EM structure determination. Nat. Methods 14, 290–296 (2017).

47. Punjani, A., Zhang, H. & Fleet, D. J. Non-uniform refinement: adaptive regularization improves single-particle cryo-EM reconstruction. Nat. Methods 17, 1214–1221 (2020).

48. Casañal, A., Lohkamp, B. & Emsley, P. Current developments in Coot for macromolecular model building of Electron Cryo-microscopy and Crystallographic Data. Protein Sci. Publ. Protein Soc. 29, 1069–1078 (2020).

49. Adams, P. D. et al. PHENIX: a comprehensive Python-based system for macromolecular structure solution. Acta Crystallogr. D Biol. Crystallogr. 66, 213–221 (2010).

50. Williams, C. J. et al. MolProbity: More and better reference data for improved all-atom structure validation. Protein Sci. Publ. Protein Soc. 27, 293–315 (2018).

51. Smart O.S., et al. Grade, version 1.2.20. Global Phasing Ltd (2021).

52. Klesse, G., Rao, S., Sansom, M. S. P. & Tucker, S. J. CHAP: A Versatile Tool for the Structural and Functional Annotation of Ion Channel Pores. J. Mol. Biol. 431, 3353–3365 (2019).

53. Pettersen, E. F. et al. UCSF ChimeraX: Structure visualization for researchers, educators, and developers. Protein Sci. Publ. Protein Soc. 30, 70–82 (2021).

54. Jo, S., Kim, T., Iyer, V. G. & Im, W. CHARMM-GUI: a web-based graphical user interface for CHARMM. J. Comput. Chem. 29, 1859–1865 (2008).

55. Huang, J. et al. CHARMM36m: an improved force field for folded and intrinsically disordered proteins. Nat. Methods 14, 71–73 (2017).

56. Vanommeslaeghe, K. et al. CHARMM general force field: A force field for drug-like molecules compatible with the CHARMM all-atom additive biological force fields. J. Comput. Chem. 31, 671–690 (2010).

57. Liu, H., Fu, H., Chipot, C., Shao, X. & Cai, W. Accuracy of Alternate Nonpolarizable Force Fields for the Determination of Protein-Ligand Binding Affinities Dominated by Cation-π Interactions. J. Chem. Theory Comput. 17, 3908–3915 (2021).

58. Abraham, M. J. et al. GROMACS: High performance molecular simulations through multi-level parallelism from laptops to supercomputers. SoftwareX 1–2, 19–25 (2015).

59. Bussi, G., Donadio, D. & Parrinello, M. Canonical sampling through velocity rescaling. J. Chem. Phys. 126, 014101 (2007).

60. Parrinello, M. & Rahman, A. Crystal Structure and Pair Potentials: A Molecular-Dynamics Study. Phys. Rev. Lett. 45, 1196–1199 (1980).

61. Hess, B. P-LINCS: A Parallel Linear Constraint Solver for Molecular Simulation. J. Chem. Theory Comput. 4, 116–122 (2008).

62. Essmann, U. et al. A smooth particle mesh Ewald method. J. Chem. Phys. 103, 8577–8593 (1995).

63. Gowers, R. et al. MDAnalysis: A Python Package for the Rapid Analysis of Molecular Dynamics Simulations. in 98–105 (Austin, Texas, 2016). doi:10.25080/Majora-629e541a-00e.

64. Hunter, J. D. Matplotlib: A 2D Graphics Environment. Comput. Sci. Eng. 9, 90–95 (2007).

65. Humphrey, W., Dalke, A. & Schulten, K. VMD: visual molecular dynamics. J. Mol. Graph. 14, 33–8, 27–28 (1996).

