## Supplementary Figures for "Cryo-EM structures of ρ1 GABA_A_ receptors with antagonist and agonist drugs"

**Supplementary Information**

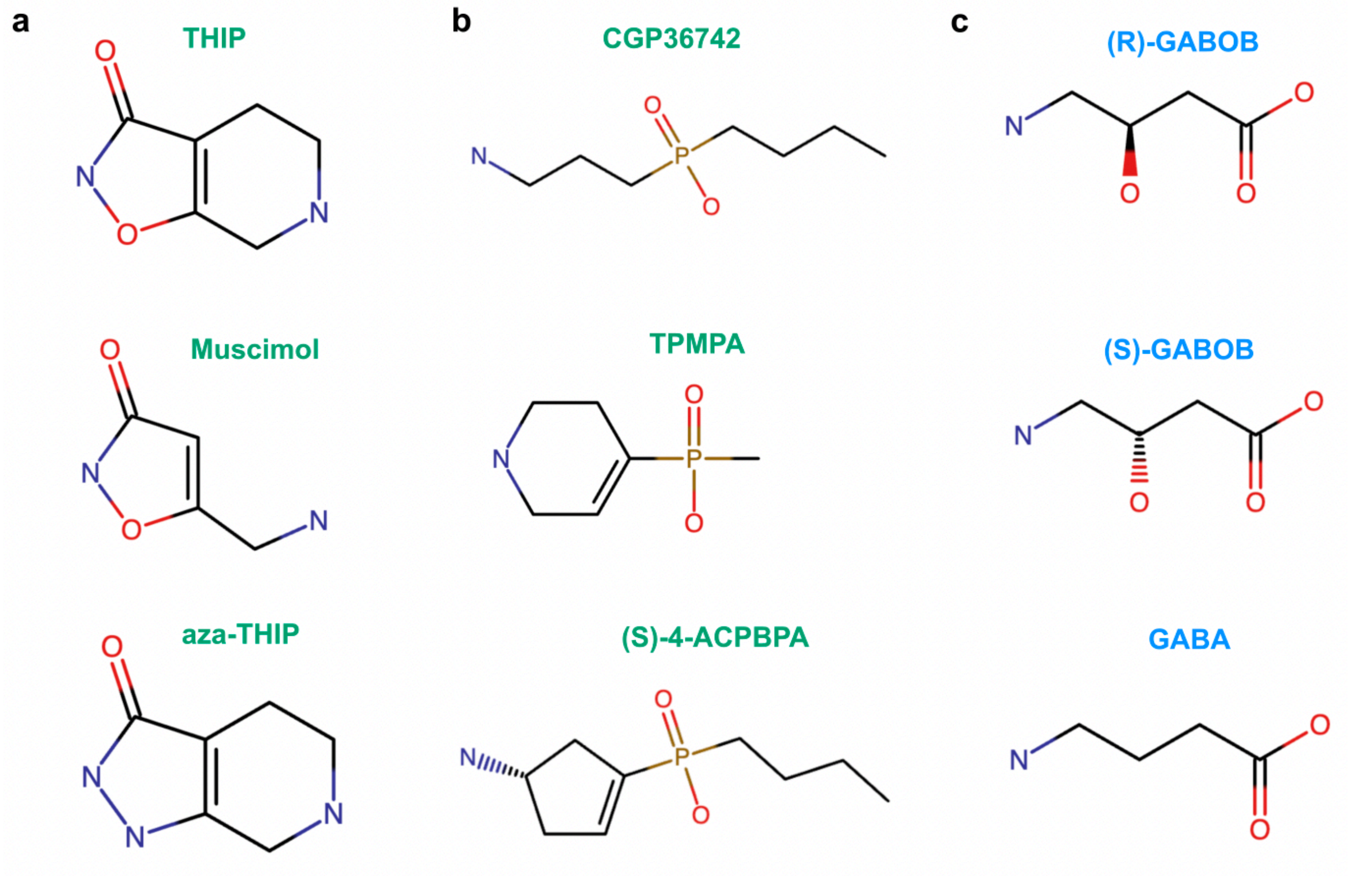


**Supplementary Fig 1. Chemical structures of relevant compounds.**

(a) Chemical structures of THIP, muscimol and aza-THIP.

(b) Chemical structures of CGP36742 and TPMPA and (S)-4-ACPBPA.

(c) Chemical structures of (R)- and (S)-GABOB and GABA.

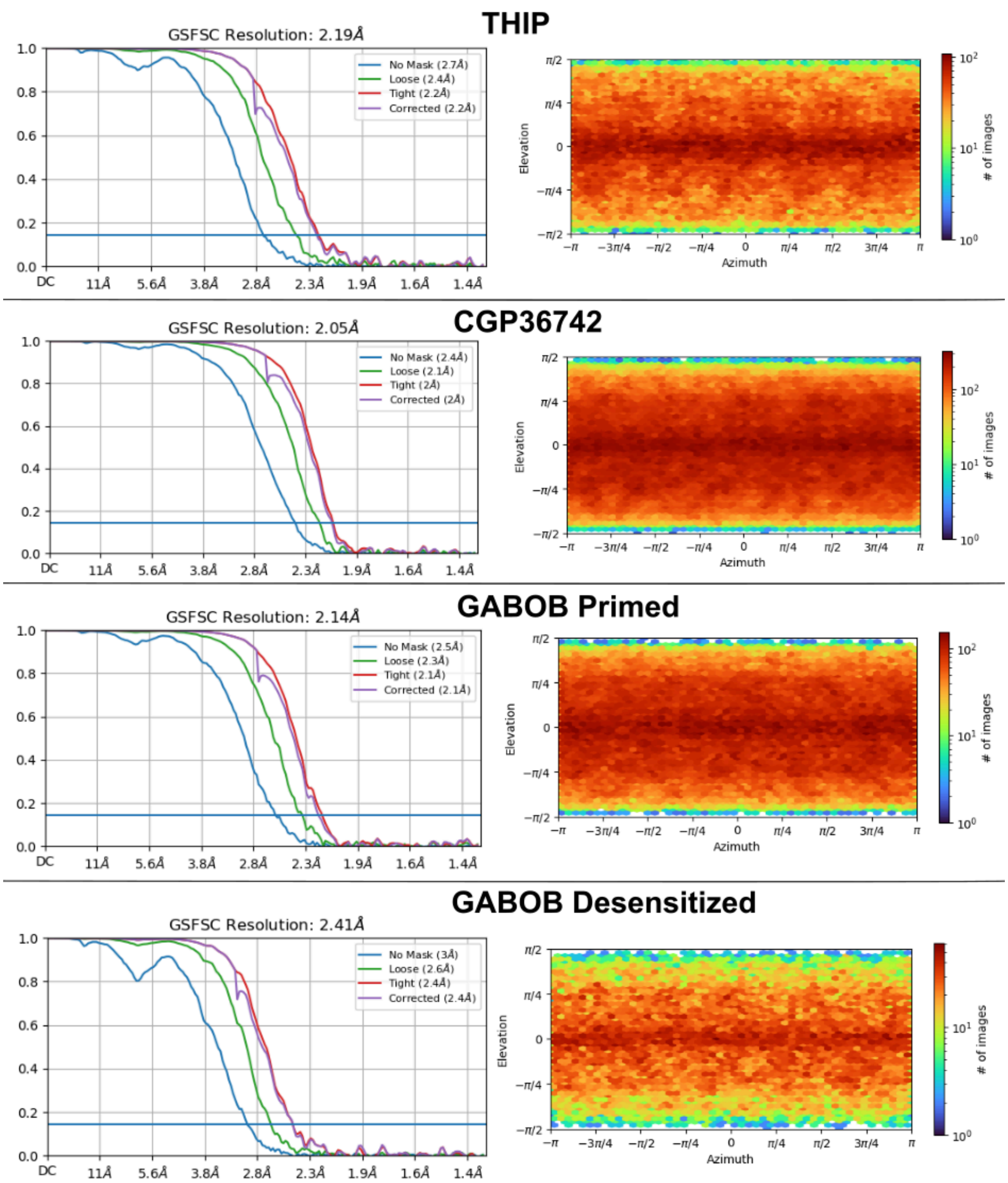


**Supplementary Fig 2. Fourier shell correlation (FSC) curves (left) and angular distributions (right) of the cryo-EM maps.**

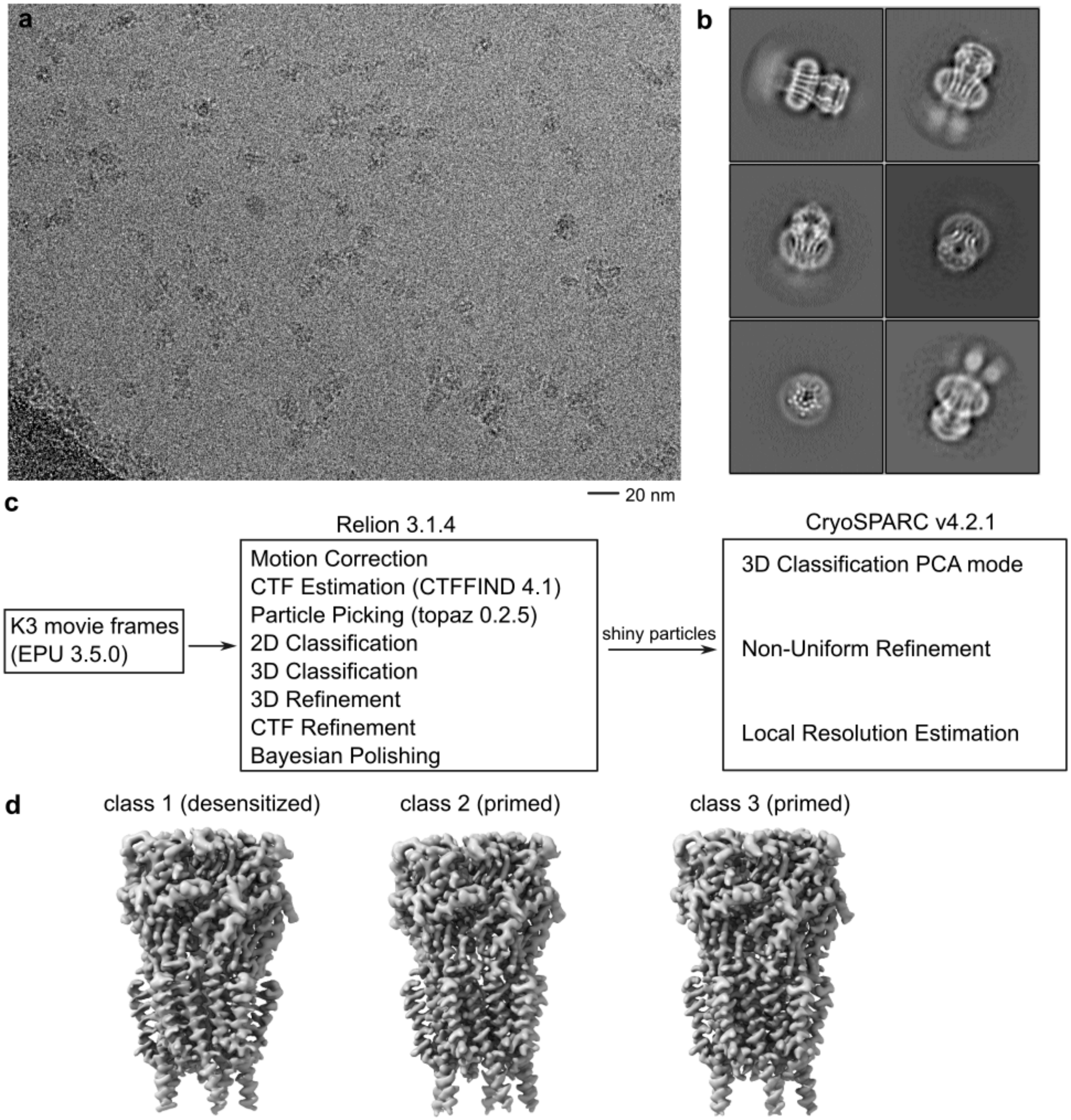


**Supplementary Fig 3. Processing pipelines for ρ1-EM structures.**

(a) Representative cryo-EM image from the ρ1-EM with CGP36742 dataset.

(b) Representative 2D classification images from the ρ1-EM with CGP36742 dataset.

(c) Cryo-EM data processing workflow for the three datasets reported in this work.

(d) Representative 3D classification reconstructions from the ρ1-EM with GABOB dataset.

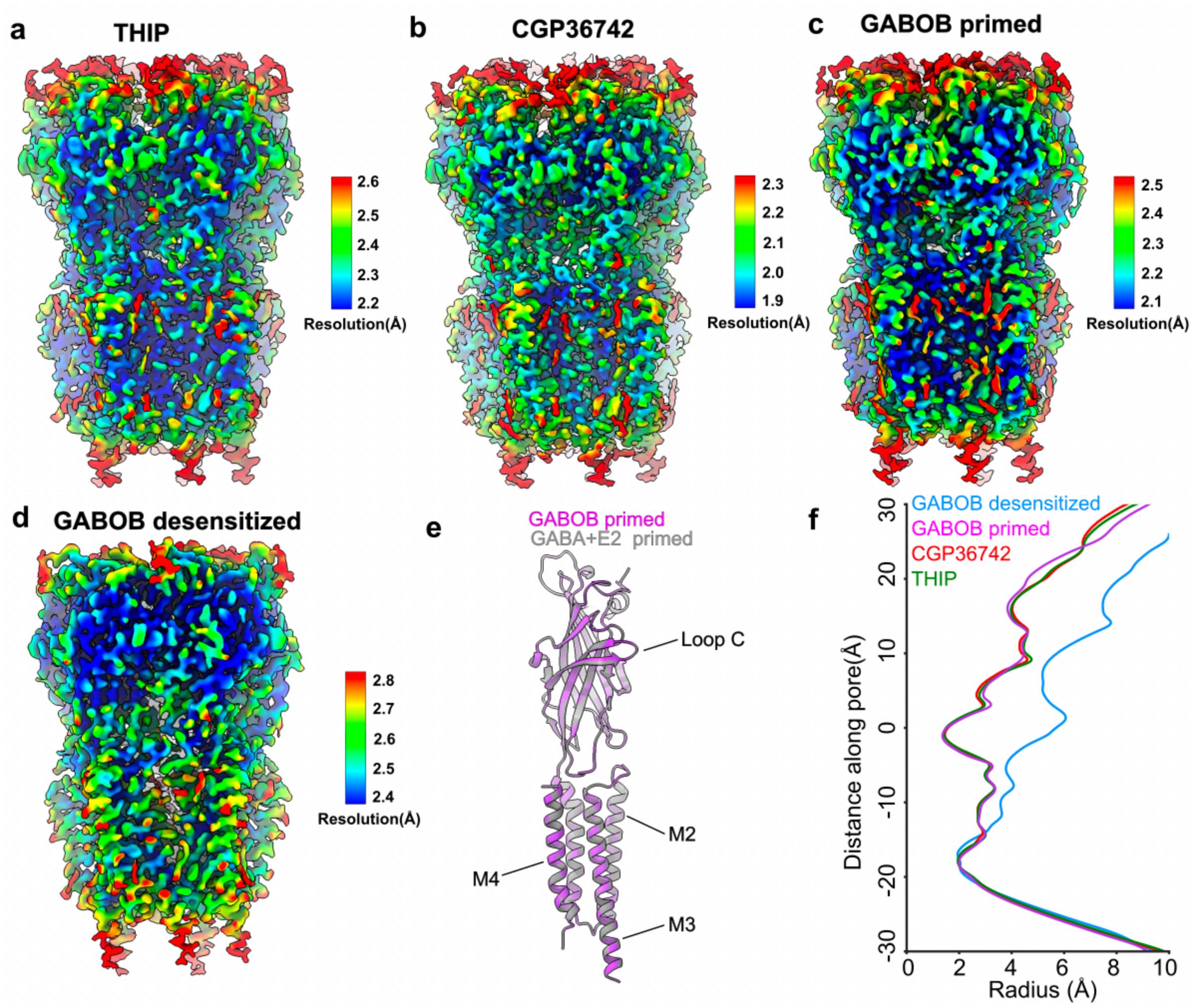


**Supplementary Fig 4. Local resolution and pore-radius profiles of the ρ1-EM structures.**

(a-d) Cryo-EM maps colored by local resolution.

(e) Superimposed structures of ρ1-EM in the primed state bound to GABOB (purple) and GABA+E2 (gray).

(f) Pore-radius profiles of ρ1-EM structures.

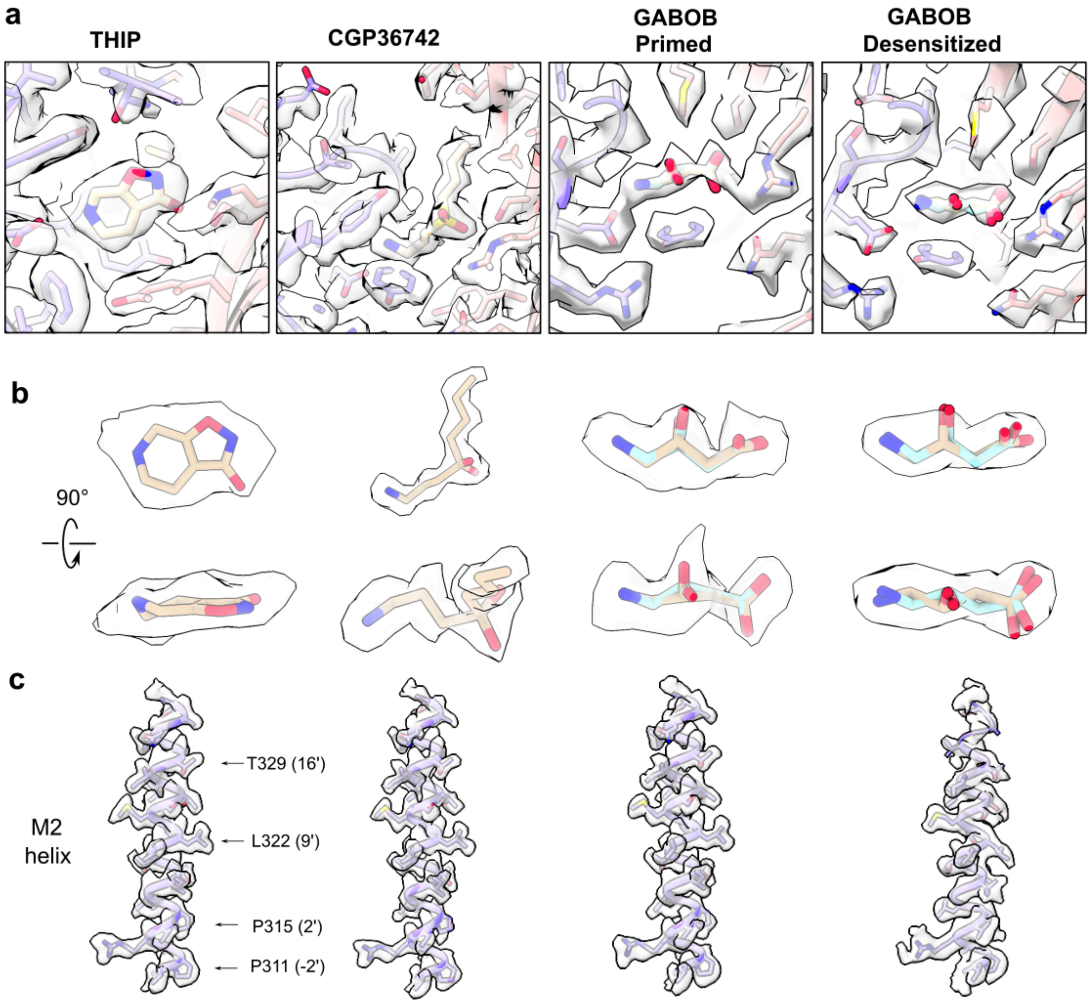


**Supplementary Fig 5. Representative densities of ρ1-EM structures**

(a) Densities and models of drug binding sites from the structures reported in this study.

(b) Densities and models of the ligands in two viewing angles.

(c) Densities and models of M2 helices from the structures reported in this study.

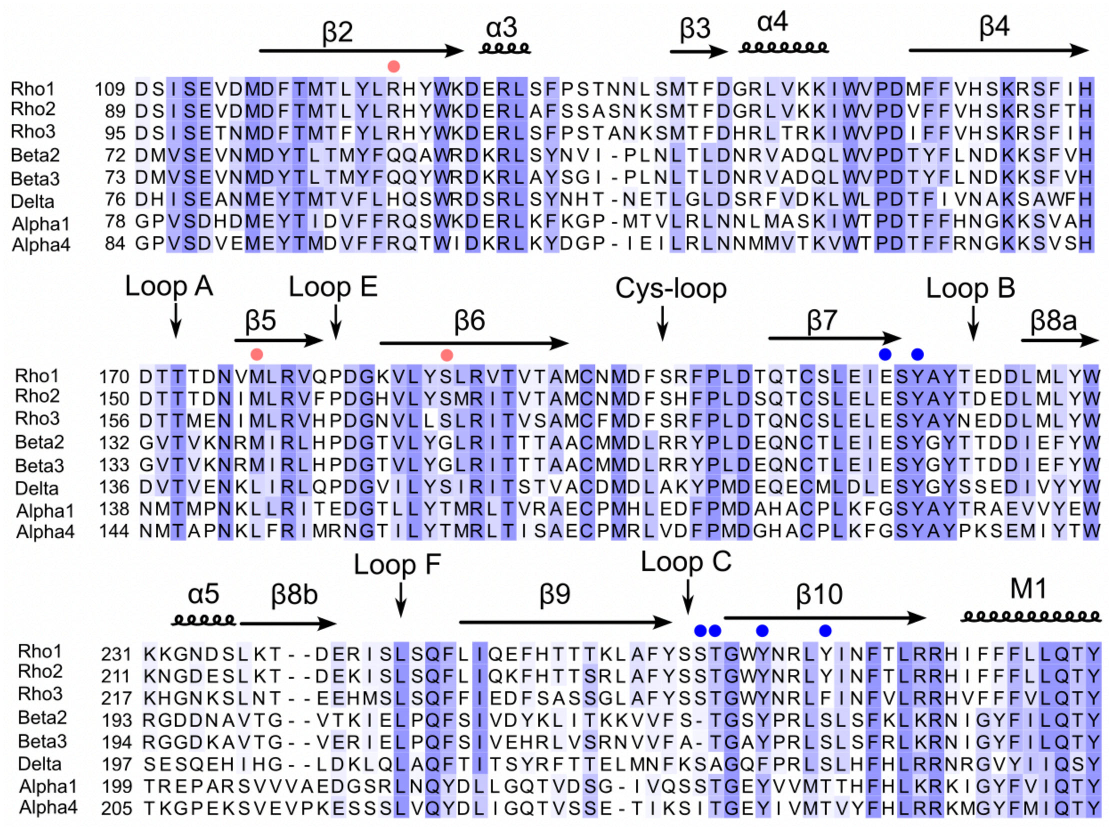

**Supplementary Fig 6. Sequence alignment of the orthosteric ligand binding region of representative human GABA$_A$ receptors.** Residues are numbered according to reference UniProt sequences, with key structural features labeled above. Dots indicate positions involved in ligand binding from the principal (blue) and complementary (red) faces.

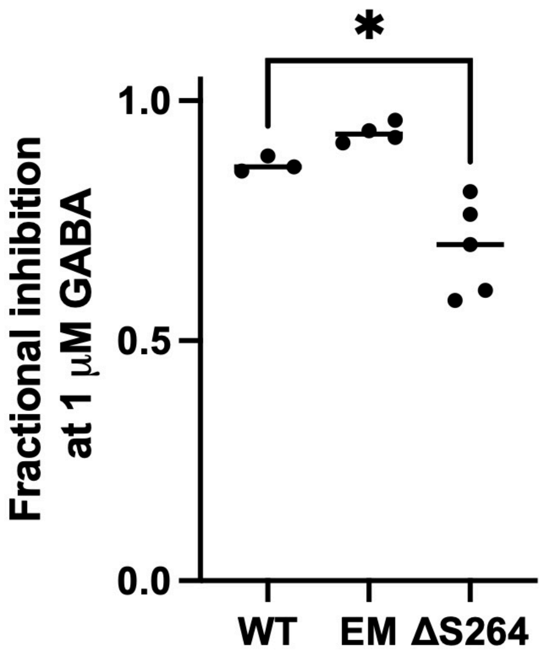


**Supplementary Fig 7. Electrophysiology profiles of ρ1 constructs**

Comparison of inhibition of 1 μM GABA response by THIP for three ρ1 constructs. Asterisk indicates significance of $p < 0.05$ in a two-way $t$-test between wild-type and ΔS264 variants of full-length ρ1 ($p = 0.0151$).

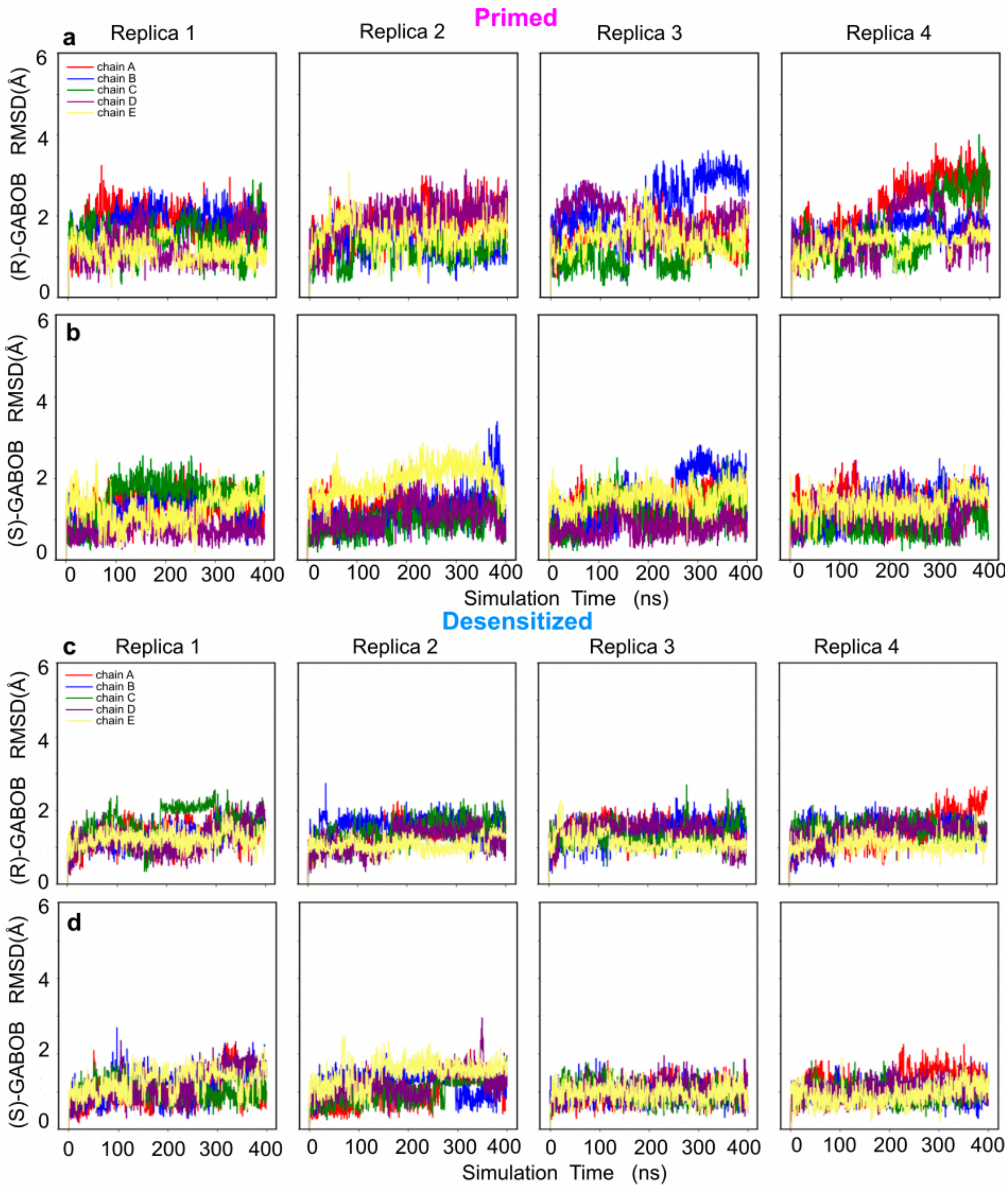


**Supplementary Fig 8. GABOB stability during MD simulations.**

(a-b) Dynamics of (R)- (above) and (S)-GABOB (below) in the primed state of ρ1-EM, calculated by RMSD and colored by chain.

(c-d) Dynamics of (R)- (above) and (S)-GABOB (below) in the desensitized state of ρ1-EM.
